# Tissue-specific clonal selection and differentiation of CD4⁺ T cells during infection

**DOI:** 10.1101/2025.08.25.672130

**Authors:** Roham Parsa, Helder Assis, Tiago B.R. de Castro, Gabriella Lima dos Reis, Arpita Sushil, Harald Hartweger, Angelina M. Bilate, Daniel Mucida

## Abstract

Pathogen-specific CD4⁺ T cells undergo dynamic expansion and contraction during infection, ultimately generating memory clones that shape the subsequent immune responses. However, the influence of distinct tissue environments on the differentiation and clonal selection of polyclonal T cells remains unclear, primarily because of the technical challenges in tracking these cells in vivo. To address this question, we generated Tracking Recently Activated Cell Kinetics (TRACK) mice, a dual-recombinase fate-mapping system that enables precise spatial and temporal labeling of recently activated CD4⁺ T cells. Using TRACK mice during influenza infection, we observed organ-specific clonal selection and transcriptional differentiation in the lungs, mediastinal lymph nodes (medLNs), and spleen. T cell receptor (TCR) sequencing revealed that local antigenic landscapes and clonal identity shape repertoire diversity, resulting in a low clonal overlap between tissues during acute infection. During the effector phase, spleen-derived CD4⁺ T cells preferentially adopted a stem-like migratory phenotype, whereas those activated in the medLNs predominantly differentiated into T follicular helper (Tfh) cells. Memory formation was associated with increased clonal overlap between lung and medLN-derived cells, whereas splenic clones retained a distinct repertoire. Additionally, memory CD4⁺ T cells displayed converging antigen specificity across tissues over time. These results highlight the tissue-dependent mechanisms driving clonal selection and functional specialization during infection and underscore how memory development facilitates clonal redistribution and functional convergence.

## Introduction

T cells play an essential role in orchestrating adaptive immune responses by differentiating into specialized effector subsets that are critical for host resistance and immune tolerance^1–4^. Extensive research has detailed the transcriptional and functional properties of these subsets and their phenotypes from initial activation in secondary lymphoid tissue, differentiation, and migration towards the target tissue^4–7^. While conventional views of T cell priming are centered around draining lymph nodes (LN), recent studies have demonstrated a role for migratory dendritic cells (mDCs) in priming influenza-specific CD8^+^ T cells within the spleen, resulting in distinct migratory patterns from LN DCs^8^. Furthermore, studies from the past two decades have indicated that lymphoid- and tissue-specific environmental signals significantly influence T cell differentiation and function^9–16^. For instance, influenza infection in humans has been shown to induce compartmentalized memory T cell responses in lymphoid versus mucosal tissues, with memory CD4^+^ T cells acquiring a different phenotype than that of CD8+ T cells^17^. Differences in the circulating patterns of memory CD4^+^ and CD8^+^ T cells have also been reported in murine models of viral infections^18^. However, the precise mechanisms underlying tissue-specific imprinting and its long-term effects on T cell function and memory generation remain poorly understood.

Another significant aspect of T cell biology that is limited by existing tools is the understanding of infection-induced T cell clonal expansion and distribution within polyclonal populations^19^. Traditional approaches to assess pathogen-specific T cells, including TCR transgenic models and MHC tetramer staining, have provided valuable insights into clonal dynamics but are constrained by their reliance on predefined epitopes, affinity and limited antigen specificities^20–23^. Recent advances in sequencing technologies, mouse genetics, and repertoire analysis have allowed comprehensive and unbiased profiling of TCR repertoires, providing an initial understanding of endogenous T cell dynamics during infections^19,24–26^. Despite these advances, the relationship between anatomical context and repertoire selection during pathogen-induced immune responses in polyclonal systems remains to be elucidated.

These gaps raise significant questions regarding the impact of priming microenvironments on the resultant T cell clonal repertoire and their roles in protective and pathogenic outcomes. To address these limitations, we developed the Tracking Recently Activated Cell Kinetics (TRACK) system, a dualrecombinase-based fate-mapping approach that enables unbiased temporal labeling of recently activated polyclonal CD4⁺ T cells, independent of predefined antigen specificity and affinity. By marking T cells transiently expressing CD62L and the activation marker CD69, the TRACK model provides a robust platform for investigating the spatial dynamics and clonal evolution of “perturbation-specific” CD4⁺ T cell responses. We employed the TRACK system to dissect how influenza virus infection shapes the transcriptional programming, clonal distribution, and functional diversity of CD4⁺ T cells across distinct anatomical sites. By leveraging single-cell RNA sequencing and high-resolution TCR repertoire analysis, our approach revealed unexpected insights into how tissue-specific priming environments influence the clonal and functional architecture of immune response.

## TRACKing recently activated T cells as a strategy to analyze pathogen-specific responses

Naïve T cells express high levels of CD62L, encoded by *Sell*, but lack the activation marker CD69. Upon activation, naïve T cells rapidly upregulate CD69 on the cell surface, resulting in the transient co-expression of CD62L and CD69, followed by the downregulation of both *Sell* and *CdCS*. Based on this expression pattern, we designed an approach to track early T cell activation during priming in mice. We generated a strain expressing CreER recombinase after the last exon of the *CdCS* gene and another strain expressing DreER after the last exon of the *Sell* gene, both of which are inducible by tamoxifen administration (Extended Data Fig. 1a). Cre recombinase excises genetic regions flanked by loxP elements, whereas Dre recombinase excises genetic regions flanked by rox elements. To generate the TRACK system, these strains were crossed to Ai66 reporter harboring a CAG-roxSTOProx-loxpSTOPloxp-tdTomato cassette in the ubiquitously expressed *Rosa2C* locus (Fig. 1a and Extended Data Fig. 1a). Tamoxifen treatment induces the excision of these two STOP cassettes that otherwise prevented tdTomato (Tomato) expression, allowing fluorescent Tomato to be permanently expressed specifically in T cells that simultaneously express CD62L and CD69 during the tamoxifen administration (Fig. 1a). Consequently, naïve, circulating, and resident memory T cells that do not co-express both markers remain unlabeled (Fig. 1a). Hence, this system presumably enables the specific labeling of recently activated T cells within a defined time window determined by tamoxifen treatment.

**Fig. 1:**
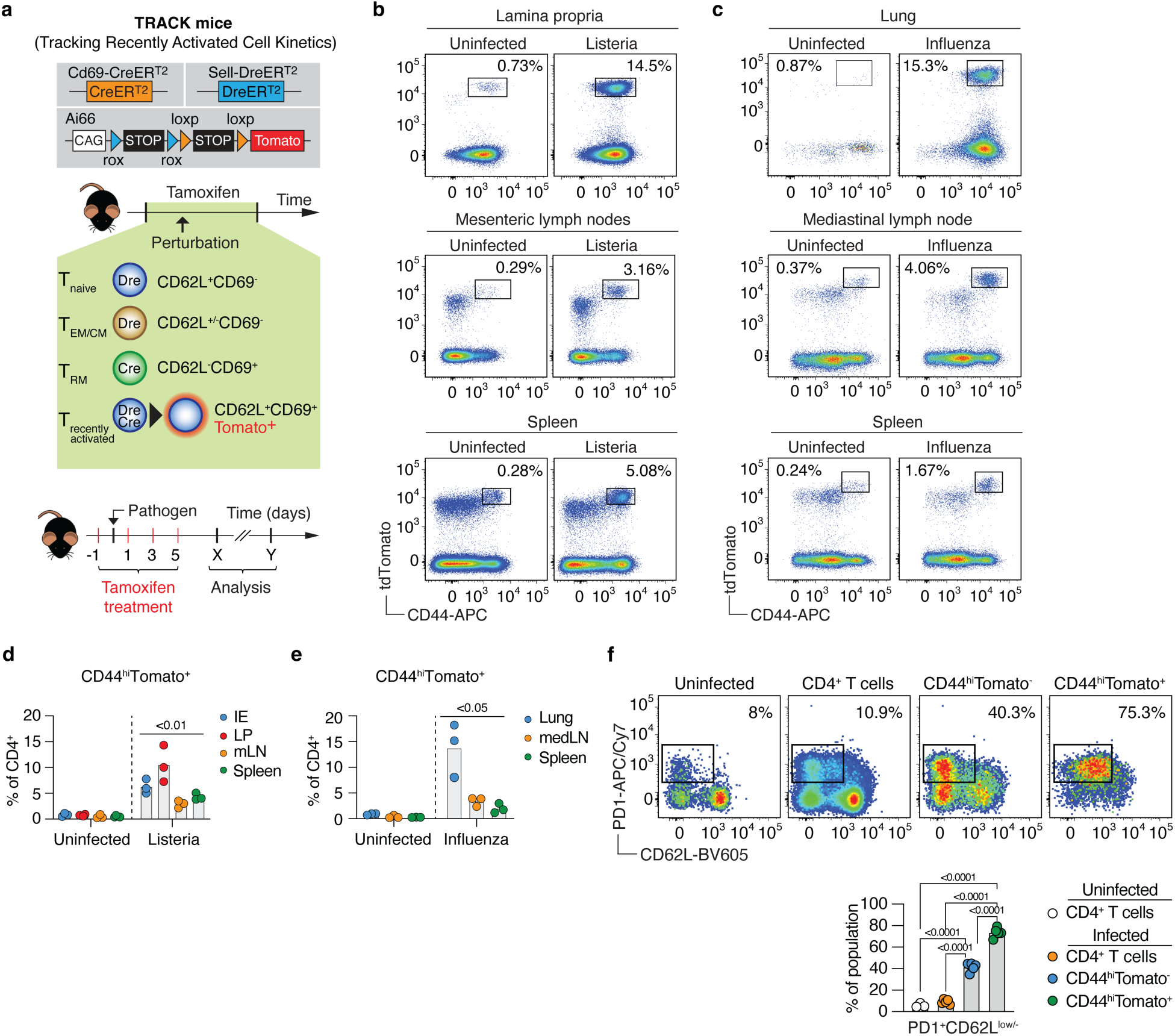
**Tracking of pathogen-specific CD4^+^ T cells during intestinal and respiratory infection.** a, The development and overview of the TRACK mice. The CreER casette was inserted after the last exon of the Cd69 gene and the DreER was inserted after the last exon of the Sell gene. TRACK mice were treated with tamoxifen at day -1, 1,3 and 5 relative to the day of infection. Tamoxifen-treated TRACK mice were infected with Listeria monocytogenes (b, d) or with influenza PR8 (c, e) and CD4^+^ T cells were analyzed 9 days post infection. Representative plots of minimum 3 independent experiments. Infected and control are compared by two-tailed Student’s t-test. f, The frequence of PD1^+^CD62L^|OW/^ in the medLN 9 days post influenza PR8 infection. Representative plot of minimum 3 independent experiments. One-way analysis of variance (ANOVA) with Tukey’s multiple-comparison test.

To assess the sensitivity of our model, we orally infected tamoxifen-treated TRACK mice with the intestinal pathogen *Listeria monocytogenes* and quantified the fate-mapped Tomato-positive CD4⁺ T cells in the intestinal epithelium (IE), intestinal lamina propria (LP), mesenteric lymph nodes (mLNs), and spleen nine days post-infection (Extended Data Fig. 1b and Fig. 1a). We observed a significant increase in activated CD44^+^ Tomato⁺ CD4⁺ T cells in the intestinal tissue, mLNs, and spleen compared to uninfected tamoxifen-treated TRACK mice (Fig. 1b,d). A similar pattern was observed for CD8⁺ T cells (Extended Data Fig. 1c).

Optimal labeling efficiency in TRACK mice was achieved after three to four tamoxifen administrations (Extended Data Fig. 1d). As expected, no fate-mapped Tomato⁺ CD4⁺ T cells were detected in the intestine when mice expressed only CreER or DreER under their respective promoters or in TRACK mice that were not treated with tamoxifen (Extended Data Fig. 1e). Infection of tamoxifen-treated TRACK mice with the mouse-adapted influenza A/PR/8/34 (PR8) virus also resulted in a significant increase in activated CD44^+^ Tomato⁺ CD4⁺ T cells in the lungs, draining mediastinal LNs (medLNs), and spleen compared to uninfected tamoxifen-treated TRACK mice (Fig. 1c,e). Within the CD44⁺Tomato⁺ population, the majority of CD4⁺ T cells exhibited high levels of the activation marker PD-1 and low levels of CD62L, indicative of an enriched population of activated effector T cells (Fig. 1f).

To determine whether cytokines can induce “bystander activation” and fate-mapping through CD69 expression, we isolated naïve T cells from the spleen of TRACK mice and cultured them with IFN-γ, IFN-β, or IL-2. As expected, stimulation with αCD3/αCD28 led to increased CD44 and CD69 expression, Tomato reporter activation, and robust cellular expansion (Extended Data Fig. 1f,g). In contrast, stimulation with IFN-γ, IFN-β, or IL-2 in the absence of TCR engagement did not elicit an activation phenotype based on CD69 or CD44 expression, nor did these conditions promote proliferation (Extended Data Fig. 1g). Notably, IFN-β induced low levels of Tomato expression predominantly in CD44⁻ naïve-like T cells, suggesting limited cytokine-driven bystander activation without full T cell activation (Extended Data Fig. 1g).

We also observed a distinct subset of T cells *in vivo* expressing lower levels of Tomato and CD44, along with markers associated with naïve T cells (Fig. 1b,c). Although this subset increased upon infection in young mice (4 weeks of age), no significant changes were detected in adult mice (8 and 13 weeks of age) (Extended Data Fig. 1h,i). This pattern indicates that Tomato⁺CD44⁻ T cells may originate from the thymus, as *Sell* and *CdCS* exhibit similar expression dynamics in both the thymus and periphery following TCR stimulation (Extended Data Fig. 1j). These findings suggest that the Tomato⁺CD44⁻ population likely comprises recent thymic emigrants, cytokine-stimulated naïve T cells, or a combination of both. Overall, these data demonstrate that the dual recombinase strategy in TRACK mice effectively labels recently activated T cells while enabling identification of fate-mapped cells across organs with minimal bystander activation.

Next, we sought to determine the efficiency of fate-mapping in TRACK mice by infecting them with either *Listeria monocytogenes* or influenza PR8 and assessing the proportion of activated pathogen-specific T cells that expressed Tomato as a proxy for dual recombinase efficiency. To identify pathogen-specific CD4⁺ T cells, we used major histocompatibility complex class II (MHC II) tetramers. *Listeria monocytogenes*-specific CD4⁺ T cells were detected using Listeriolysin O peptide (LLO₁₉₀₋₂₀₁) tetramers, whereas influenza PR8-specific CD4⁺ T cells were identified using nucleoprotein peptide (NP₃₁₁₋₃₂₅) tetramers. On average, 43 % of tetramer-binding pathogen-specific CD4⁺ T cells expressed Tomato (Fig. 2a,b). This observation highlights the inherent limitations of the dual recombinase-mediated fate-mapping strategy, while demonstrating that TRACK mice provide a reliable tool for capturing a substantial fraction of recently activated pathogen-specific T cells.

**Fig. 2:**
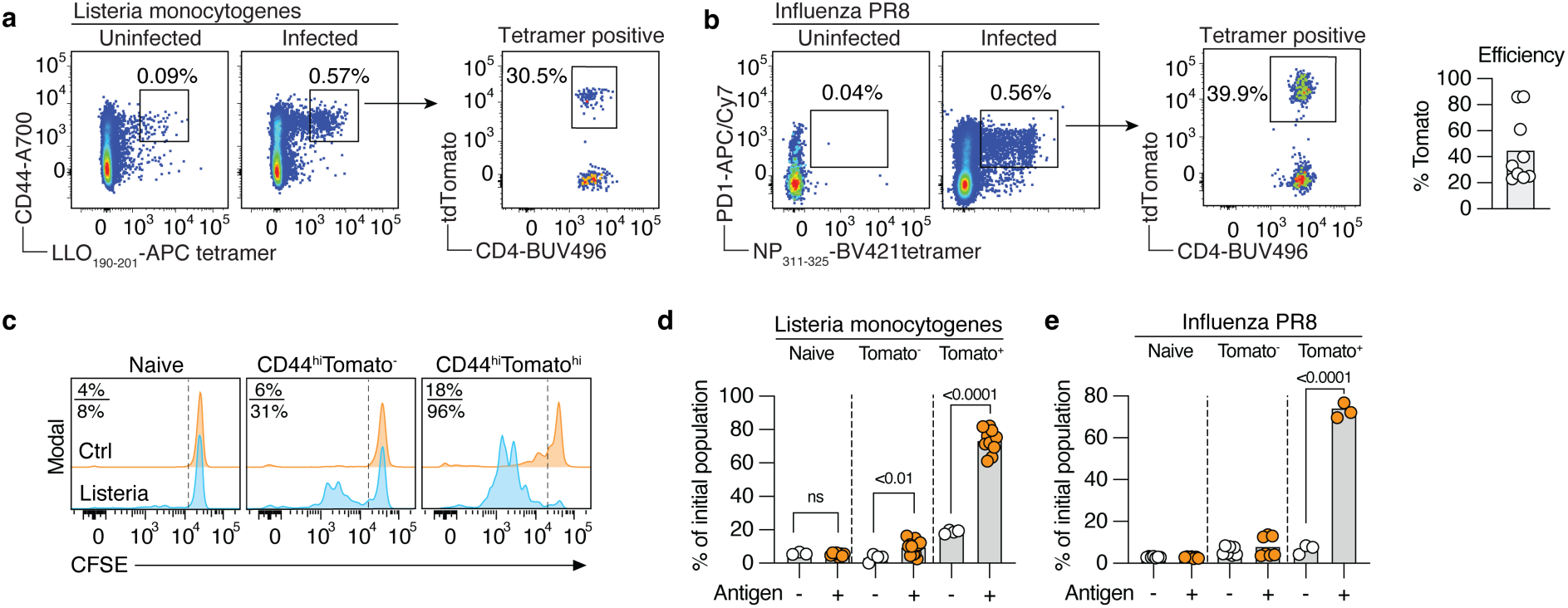
**Enrichment of perturbation-specific T cells.** a, The frequency of Tomato^+^ among LLO_190 201_-specific CD4^+^ T cells in mesenteric lymph nodes 9 days post Listeria infection, b, The frequency of Tomato^+^ among NP_311 325_-specific CD4^+^ T cells in medLN 9 days post influenza infection. Right plot shows frequences of Tomato^+^CD4^+^ T cells pooled from both Listeria and influenza infected mice. Two independent experiments. CD44^|OW^CD62L^+^Tomato’ (naive), CD44^h^Tomato’ and CD44^h^Tomato^+^ CD4^+^ T cells were isolated from the spleen of L. monocytogenes infected TRACK mice 30 days post infection (c, d) or influenza PR8 infected medLN and spleen (e) 30 days post infection. T cells were stained with CFSE and co-cultured for 48-72 hours with splenic DCs that were pulsed with heat-killed pathogen. Pooled data from minimum 2 experiments, n = 3-10. Statistical differences were determined with two-tailed Student’s t-test.

Finally, we determined the proportion of fate-mapped T cells that specifically responded to pathogens. We infected tamoxifen-treated TRACK mice with *Listeria monocytogenes* and, 30 days post-infection, isolated naïve (CD44⁻/^low^CD62L⁺Tomato⁻), and “putative recently activated” CD44⁺Tomato⁻ or CD44⁺Tomato⁺ CD4⁺ T cells from the spleen. Isolated T cells were stained with carboxyfluorescein succinimidyl ester (CFSE), co-cultured with DCs, and stimulated with heat-killed *Listeria monocytogenes (Lm)*. To assess the fraction of antigen-responsive cells, we measured the CFSE dilution as an indicator of cell division. No CFSE dilution was observed in naïve T cells stimulated with heat-killed *Listeria monocytogenes*, but approximately 30% of CD44⁺Tomato⁻ divided, likely reflecting the fraction of pathogen-specific T cells escaping TRACK fate-mapping. Nevertheless, 75-85% of CD44⁺Tomato⁺ CD4⁺ T cells proliferated in response to *Lm*-pulsed DCs (Fig. 2c,d). Similar results were observed in the influenza PR8 model (Fig. 2e), confirming the robustness of our approach for different infections. These findings demonstrate that the vast majority of fate-mapped T cells identified using the TRACK model respond to pathogens, further validating the utility of this model for tracking antigen-experienced T cells after infection in both lymphoid and non-lymphoid organs.

## Divergent CD4⁺ T cell fates in lymphoid and non-lymphoid organs after influenza infection

During the early stages of influenza infection, DCs in the lungs capture pathogen antigens and migrate to draining lymph nodes. Lung-derived migratory DCs can be detected in the medLN as early as 12 h after infection. These migratory DCs can subsequently egress the medLN and reach the spleen within 24 h post-infection^27^, thereby enabling another potential site for T cell activation in response to pathogens located in tissues. However, whether T cell differentiation, functional specialization, and migration differ between these inductive sites remains unclear. Previous studies have shown that naïve CD8⁺ T cells can be activated in the spleen, independently of the medLN, within a few days after influenza PR8 infection^27–29^. To determine whether CD4⁺ T cells can also be activated independently in the spleen, we infected CD45.1 congenic C57BL/6J mice with influenza PR8 and transferred naïve NP-specific CD45.2⁺ CD4⁺ T cells into the CD45.1^+^ recipients. To prevent the migration of these T cells to the medLN while still allowing access to the spleen, mice were treated with anti-CD62L antibodies (Extended Data Fig. 2a). Anti-CD62L treatment effectively blocked the migration and activation of NP-specific CD4⁺ T cells in the medLN, but they continued to be activated in the spleen, further confirming the spleen as an independent site of CD4⁺ T cell activation during influenza PR8 infection (Extended Data Fig. 2b).

To elucidate the functional and clonal distribution of virus-specific CD4⁺ T cells, we infected tamoxifen-treated TRACK mice with influenza PR8 and isolated CD44⁺Tomato⁺ T cells from the medLN, spleen, and lungs 9- and 56-days post-infection. We then performed single-cell RNA sequencing (scRNA-seq) to obtain gene expression and T cell receptor (TCR) repertoire data. These two points represent the effector (day 9) and memory (day 56) stages of CD4⁺ T cell responses (Extended Data Fig. 2c). We identified 15 CD4^+^ T cell states across organs and timepoints (Fig. 3a,b).

**Fig. 3.**
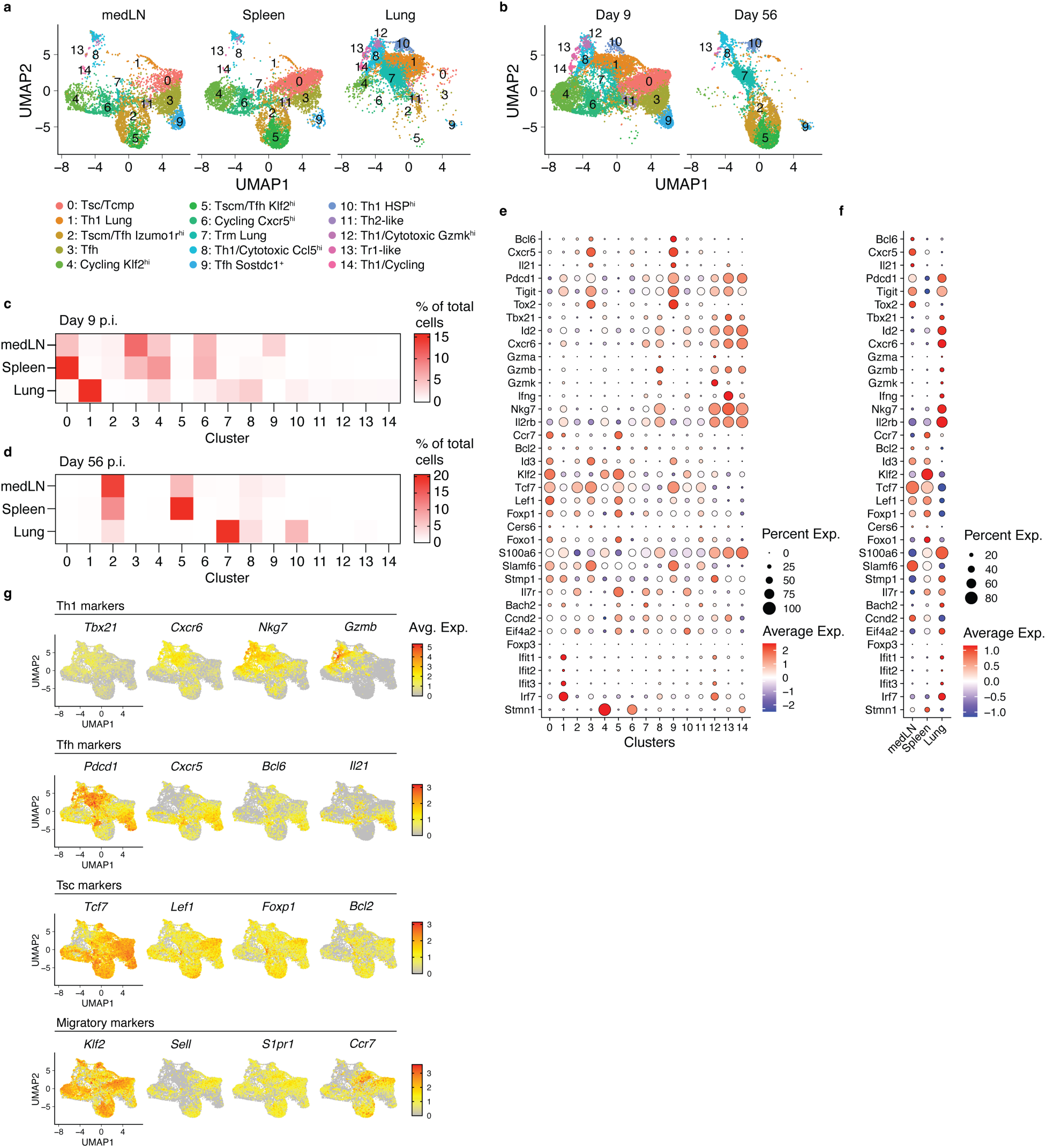
: Distinct transcriptional profiles among pathogen-specific CD4^+^ T cells across lymphoid and non-lymphoid organs TRACK mice were infected with influenza PR8 and hash-tagged CD4^+^Tomato^+^ T cells were isolated from (a) the medLN, spleen and lungs (b) at day 9 or day 56 post infection and analyzed by RNA-seq. Two seperate experiments, day 9 n=3, day 56 n=4. Percentage of cells distributed across clusters and organs at day 9 (c) and 56 (d) post-infection. Pooled data from a and b. e, Signature gene expression across clusters from day 9 and day 56 post infection, f, Average gene expression across organs from day 9 and day 56 post infection g, Key gene expression forThl *(Tbx21, Cxcr6, Nkg7, Gzmb),* Tfh *(Pdcdl, Cxcr5, Bcl6, 1121),* Tsc *(Tcf7, Left Foxpt Bcl2)* and migratory *(Klf2, Sell, S1pr1, Ccr7)* across organs and timepoints. Pooled RNAseq data from 3 mice (day 9) or 4 mice (day 56) in all figures. All presented genes have adj. p-value < 0.001.

As expected, fate-mapped T cells in the lungs exhibited a distinct gene expression profile compared to those in secondary lymphoid organs (SLOs) during both the effector and memory phases (Fig. 3a,b and Extended Data Fig. 2d,e). At the effector stage, a large fraction of CD4⁺ T cells in the lungs displayed a T helper 1 (Th1) and cytotoxic phenotype, found in clusters 1 and 8, characterized by the expression of *Tbx21*, *Ifng*, *Id2*, *Nkg7*, and *Gzmb* (Fig. 3a,c,e,g and Extended Data Fig. 2d,f). In contrast, the predominant CD4⁺ T cell subsets in SLOs consisted of T follicular helper (Tfh; clusters 3 and 9) and stem-like T (Tsc; cluster 0) cells, defined by the expression of *Pdcd1*, *Cxcr5*, and *BclC* (Tfh, Fig. 3g,e), as well as *Tcf7*, *Lef1*, *Foxp1*, and *Bcl2* (Tsc, Fig. 3g,e)^30,31^.

CD4⁺ T cells in the medLN exhibited a bias toward T follicular helper (Tfh) differentiation, whereas those in the spleen showed a preference for Tsc differentiation (Fig. 3f). We also identified three large cycling clusters across all organs. Cluster 6 expressed Tfh-associated genes and was equally found in the medLN and the spleen. Cluster 4 expressed migratory genes and was enriched within the spleen. Cluster 14 expressed Th1- and lung-associated genes and was predominantly found in the lungs.

Furthermore, analysis of the top 10 differentially expressed genes among CD4⁺ T cells in the lungs revealed the presence of lung-associated genes, including *Vps37b*, *Dennd4a* and *Dusp1* (Fig. 4a). In contrast, CD4⁺ T cells in the medLN exhibited high expression of *Cxcr5* and other genes associated with Tfh differentiation, including *Cebpa* and *Il21*. Conversely, *Klf2* was the most differentially expressed gene in CD4⁺ T cells from the spleen. To determine whether these organ-specific transcriptional programs reflect local activation or post-activation migration, we analyzed clonal distributions across organs using the Gini index as a measure of clonal evenness. A higher Gini index indicates that a clone is evenly distributed across the three organs, whereas a Gini index of 0 denotes complete segregation of that clone to a single organ. Segregated clones, those that most likely underwent activation and expansion within the organ where they are predominantly detected, displayed organ-specific functional differentiation compared to more evenly distributed clones (Extended Data Fig. 2g).

**Fig. 4:**
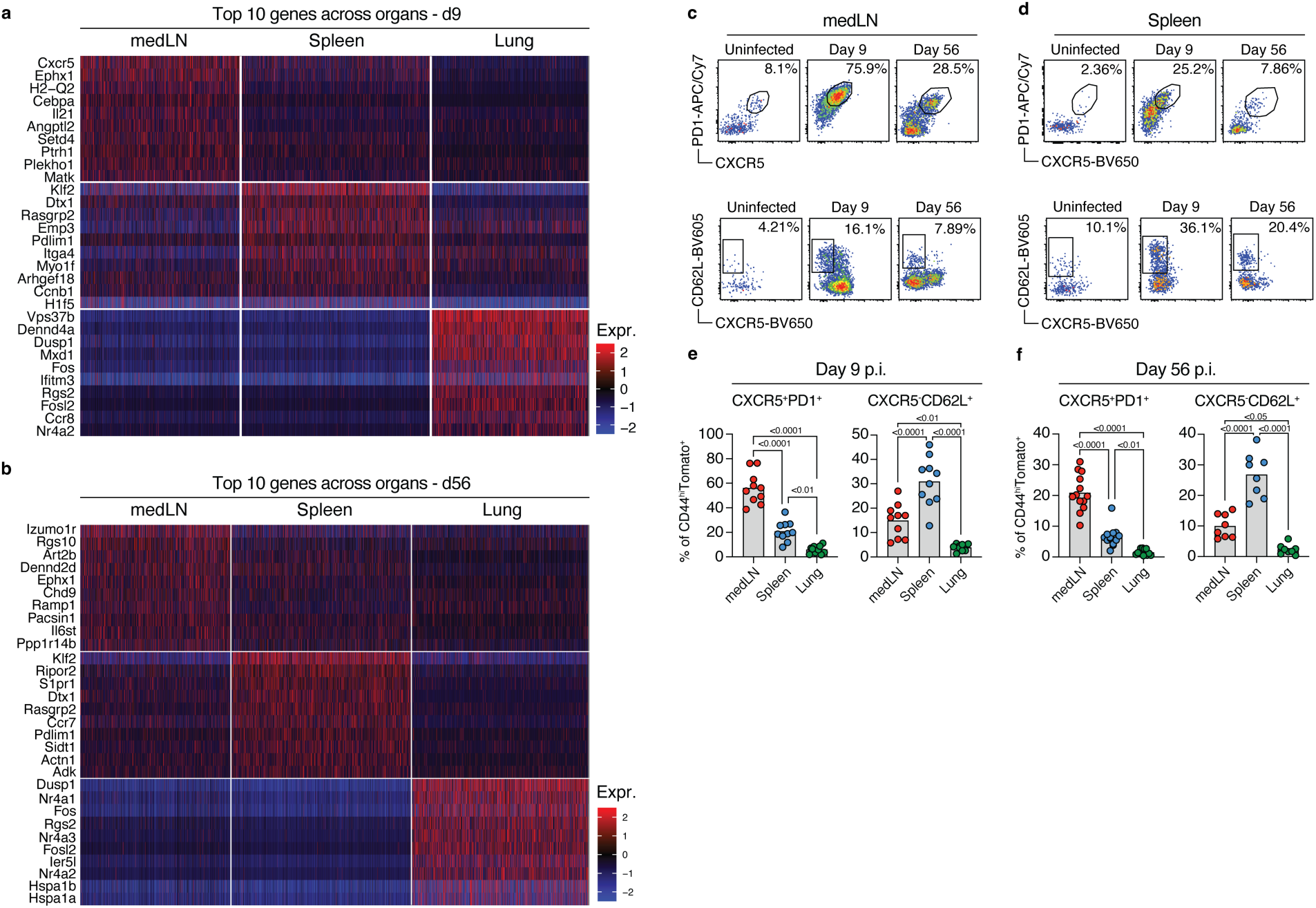
**Divergent CD4^+^ T cell functional fates in secondary lymphoid organs during influenza infection.V** The top 10 genes among all CD4^+^CD44^hl^Tomato^+^ T cells in each organ at (a) day 9 and (b) day 56 post infection. Adjusted p value < 0.0001. TRACK mice were infected with influenza PR8 virus and CD4^+^CD44^h^Tomato^+^ were analyzed for Tfh (PD1^+^, CXCR5^+^) or migratory stem-like (CD62L^+^, CXCR5) features in medLN (c) or spleen (d) at day 9 (e) or day 56 (f) post infection. Pooled data from 3 independent experiments, n=10. One-way analysis of variance (ANOVA) with Tukey’s multiple-comparison test. All presented genes have adjusted p-value < 0.0001.

At the memory stage, we identified six large CD4⁺ T cell subset clusters across organs (Fig. 3a,b,d,e and Extended Data Fig. 2e,f). Memory CD4⁺ T cells in the lungs, primarily within cluster 7, exhibited a distinct gene signature associated with the resident memory T cell (Trm) phenotype, including *CxcrC* and *Nr4a3* expression. The majority of memory CD4⁺ T cells in the medLN and spleen were distributed across two major clusters: a Tsc/Tfh-like population (cluster 2) expressing Tfh-associated genes, such as *Izumo1r*, *Cxcr5*, and *Tox*, and a Tsc/Tcm population (cluster 5) expressing key migration-related genes, including *Sell*, *S1pr1*, *Ccr7*, and *Klf2*. Both clusters expressed genes associated with stem-like properties, including *Tcf7*, *Lef1*, and *Foxp1* genes (Fig. 3g,e). *Klf2* remained highly expressed in CD4⁺ T cells in the spleen at the memory stage, whereas *Izumo1r* was the most differentially expressed gene in medLN CD4⁺ T cells (Fig. 3f and Fig. 4b). Hence, similar to the effector stage, CD4⁺ T cells in the medLN showed a bias toward the Tsc/Tfh subset, whereas those in the spleen exhibited a preference for the Tsc/central memory T cell (Tcm) subset. The *Sell* gene is a key target of the *Klf2* transcription factor, and analysis of CD62L (encoded by *Sell*), PD1, and CXCR5 surface expression in CD4⁺ T cells confirmed the distinct Tfh/Tsc bias between the medLN (Fig. 4c) and spleen (Fig. 4d) on days 9 and 56 post-infection (Fig. 4e,f). Together, these findings reveal that CD4⁺ T cells primed in different secondary lymphoid organs exhibit distinct transcriptional profiles, with medLN favoring Tfh differentiation, spleen promoting a stem-like migratory phenotype, and lungs fostering tissue-resident effector cells, highlighting organ-specific CD4⁺ T cell fate decisions during influenza infection.

## Influenza-specific CD4^+^ T cells show distinct clonal expansion and distribution across organs

Next, we sought to determine whether biased CD4⁺ T helper differentiation in the medLN and spleen influenced the seeding of CD4⁺ T cell clones in the lungs during the effector stage of influenza infection. To this end, we extracted clonal information from our scRNA-seq data, and additional TCR information was gathered by infecting tamoxifen-treated TRACK mice with influenza PR8 virus. CD44^hi^Tomato^+^CD4^+^ T cells were isolated at various time points and analyzed using scTCR-seq. We used TCR sequences as barcodes to assess clonal sharing across organs and applied the Morisita-Horn overlap index to quantify repertoire similarity, where a value of 1 indicates complete clonal overlap and a value of 0 indicates entirely distinct repertoires.

Our data revealed significantly greater clonal sharing among the expanded clones between the spleen and lungs during the effector stage than that between the medLN and lungs (Fig. 5a). To further explore the phenotypic distribution of CD4⁺ T cells across organs, we analyzed the five most expanded clones in each organ and determined their distribution across the clusters and tissues (Extended Data Fig. 3a). In the spleen, one-third of the most expanded clones were localized within clusters 0 and 4 (Fig. 5b) and were associated with migratory stem-like properties (Fig. 3e). Approximately 20 % of these expanded clones were shared with cluster 1, which was found almost exclusively in the lung. In contrast, fewer than 5 % of the five most expanded clones in the spleen were shared with the medLN at this stage of infection. To determine whether the spleen is a significant source of effector CD4^+^ T cells that migrate to the lungs, we infected splenectomized TRACK mice and determined T cell infiltration in the lungs 9 days post-influenza PR8 infection. In agreement with our previous data, splenectomized TRACK mice had fewer CD4^+^ T cells in their lungs than non-splenectomized control mice (Fig. 5c). In contrast to the spleen, over 40 % of the expanded clones in the medLN were found in clusters 0, 2, 3, and 7, all of which expressed Tfh-associated genes, such as *Cxcr5*, *BclC*, and *Il21*. Only 10 % of the most expanded medLN clones were shared with the lungs (cluster 1), and approximately 15–20 % were shared with the spleen, primarily within clusters 0, 2, 3, and 4. These findings suggest that the *Klf2*^hi^ migratory stem-like phenotype of influenza-specific CD4^+^ T cells in the spleen facilitates increased clonal sharing with the lungs, particularly in the Th1-associated cluster 1, whereas expanded clones in the medLN largely remain SLO Tfh cells. Analysis of expanded clones in the lungs suggested local clonal expansion, with fewer than 10 % of clones shared with the spleen (cluster 0) and minimal overlap with the medLNs (Extended Data Fig. 3b). Overall, this indicates that although expanded clones are shared across organs, effector CD4⁺ T cell repertoires remain largely distinct, with limited inter-organ overlap.

**Fig. 5:**
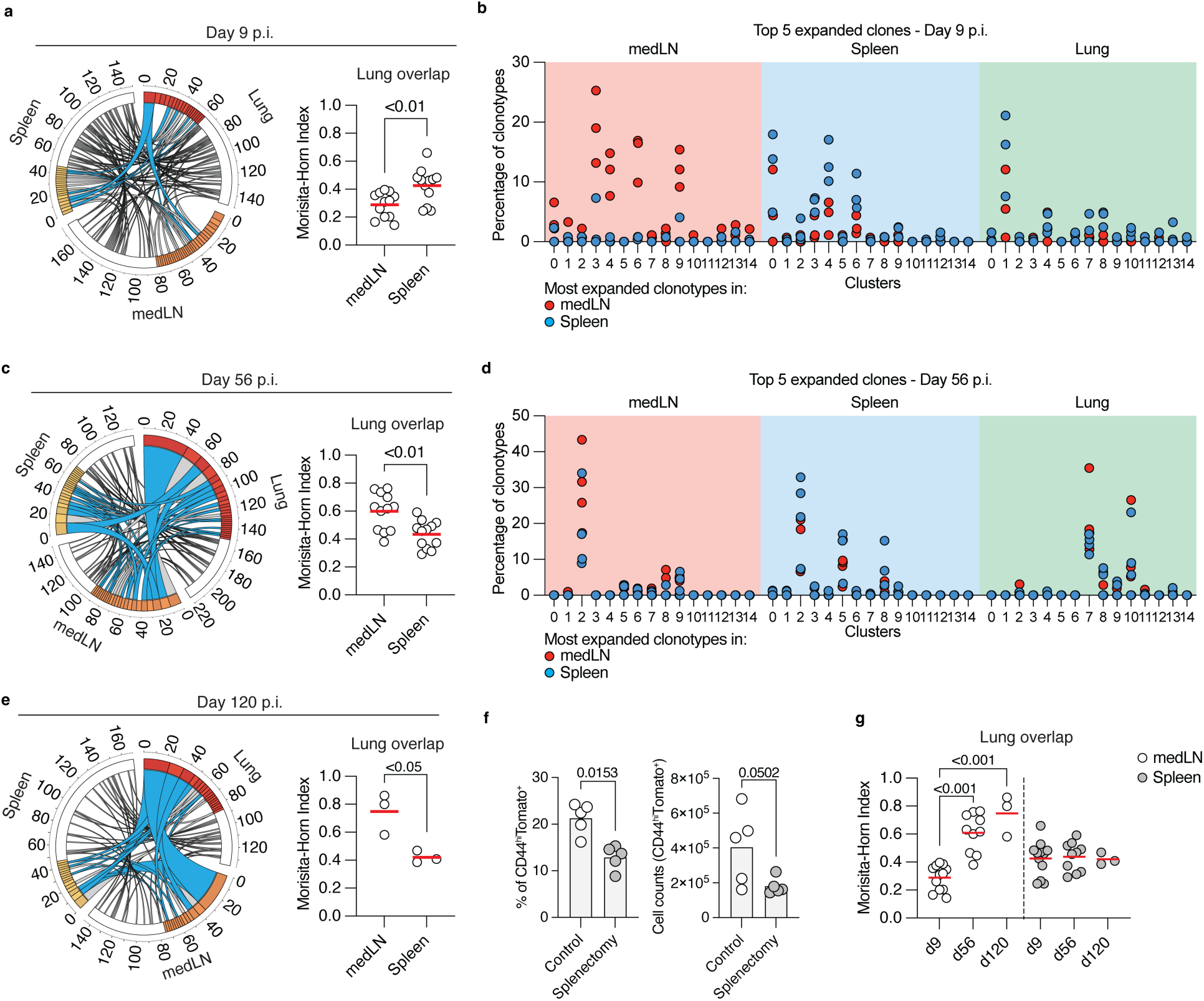
Distinct clonal expansion and distribution across organs. a, Circos plot of day 9 scTCRseq data, each segment represents organ clonal repertoire, each colored unit represents expanded clonotype, white unit represents singletons, connections along segments represents clonotype sharing across organs, colored connections represents expanded clonotypes shared. Outside ticks represtens clone counts. Representative circos plot of one mouse, pooled dot plot of 12 mice across minimum 3 independent experiments. Data from scRNAseq included, n=3. Significance among Morisita-Horn Index (MHI) data determined by Student’s T-test. b, Distribution across clusters and orgads of most expanded clones found in medLN or Spleen 9 days post infection. Each dot within cluster represents one mouse, n=3. c, Circos plot of day 56 scTCRseq and scRNAseq data. Representative circos plot of one mouse, pooled dot plot of 12 mice across minimum 3 independent experiments. Significance among MHI data determined by Student’s T-test. d, Distribution across clusters and orgads of most expanded clones found in medLN or Spleen 56 days post infection. Each dot within cluster represents one unqiue mouse, n=4. e, Circos plot of day 120 scTCRseq and scRNAseq data. Representative circos plot of one mouse, pooled dot plot of 3 mice across minimum 2 independent experiments. Significance among MHI data determined by Student’s T-test. f, Splenectomized TRACK mice were infected with influenza PR8 and CD4^+^CD44^hi^Tomato^+^ were analyzed within the lungs 9 days post infection. Representative data from one experiment n=5/group, 2 independent experiments. Significance determined by Student’s T-test. g, MHI data from (a, d and e) pooled together and analyzed with one-way analysis of variance (ANOVA) with Tukey’s multiple-comparison test.

Analysis of clonal sharing during the memory stage revealed a significant increase in overlap between the medLN and lungs at 56- and 120-days post-infection (Fig. 5d,e,g). This finding aligns with recent reports suggesting retrograde migration of Trm cells from the tissue back to the draining LNs after infection^32–34^. In contrast, the clonal overlap between the spleen and lungs of the expanded clones remained unchanged over time. Trm cells in the lungs primarily shared clones with cluster 0 (Tscm/Tfh, Izumo1r) and cluster 1 (Tsc/Tcm, Klf2) populations found in both the medLN and spleen (Fig. 5f and Extended Data Fig. 3c). These data demonstrate that over time, pathogen-specific clones become distributed throughout the host, particularly between the medLN and lungs, leading to clonal equilibrium.

Our analysis revealed that, during the effector stage, the spleen exhibited greater clonal sharing with the lungs. This observation may reflect a “division of labor” between inductive sites, in which the CD4^+^ T cells in the medLN are mostly tasked with B cell help, whereas the spleen seeds the lung with effector T cells. In contrast, clonal sharing between the medLN and lungs eventually increases during the memory phase, suggesting a gradual redistribution of tissue-resident clones to draining lymphoid organs, potentially contributing to long-term immune surveillance, broader inter-organ dissemination and recall responses.

## TCR antigen specificity shapes organ-specific clonal repertoires

Our analysis during the effector stage revealed that while some expanded CD4⁺ T cell clones were shared across organs, the overall organ repertoires remained largely distinct, suggesting that organ-specific factors influence T cell selection. One possible explanation is that antigen availability and presentation in each organ drive the selection and expansion of specific T cell clones based on TCR specificity. To investigate this, we cloned the TCRs of the most expanded clonotypes from each organ and assessed their antigen specificities (Supplementary Table 1). The cloned TCRs were stably transduced into T cell hybridomas lacking endogenous TCR expression, and successful TCR expression was confirmed using flow cytometry. TCR activation in the hybridomas was detected via GFP expression driven by the NFAT promoter, where GFP levels correlated with the TCR signal strength (Extended Data Fig. 4a). Of the 15 cloned TCRs from the effector stage, 14 showed productive TCR expression by the T cell hybridomas (Extended Data Fig. 4b).

TCR-expressing hybridomas were co-cultured with DCs pulsed with either whole viral particles or individual viral proteins, and TCR specificity was determined based on green fluorescent protein (GFP) expression. All tested clones exhibited some degree of reactivity to the viral particles (Extended Data Fig. 4c). Four of the five TCRs derived from the most expanded clones in the spleen demonstrated high reactivity against the viral particles (Fig. 6a and Extended Data Fig. 4c). In particular, the two largest clonotypes in the spleen were reactive to hemagglutinin (HA), one of the most abundant proteins in the virus^35,36^. These highly expanded HA-reactive clones were enriched in migratory stem-like clusters 0 and 4, as defined by *Klf2* and *Tcf7* expression. Although these top expanded spleen-derived clones were also shared with the lungs, they were not among the most expanded clones (Fig. 6a). In contrast, the most expanded clonotype in the lung was reactive to the polymerase subunit PB1, a less abundant viral protein highly expressed in infected epithelial cells. Similarly, the second most expanded lung clonotype responded to nucleoprotein (NP), a protein that is highly expressed in infected cells and abundant in viral particles. These dominant lung clonotypes were primarily found in Th1 cells and cytotoxic clusters 1, 6, and 8 of the lung. Finally, three of the four cloned TCRs from medLN exhibited strong reactivity to NP, with two of these clonotypes being enriched in Tfh clusters 2 and 7, respectively.

**Fig. 6.**
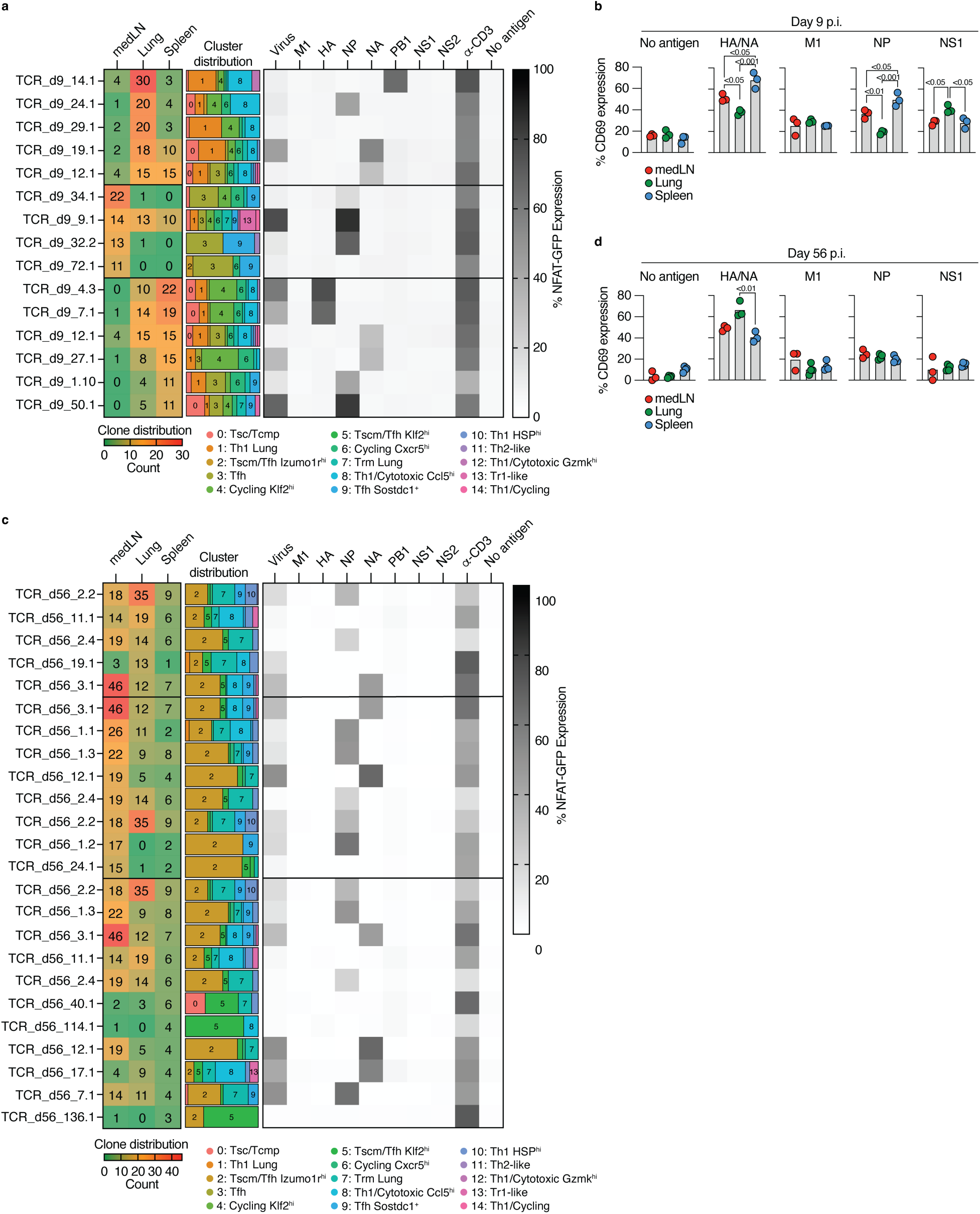
: **Clonal selection is shaped by organ-specific antigenic landscapes.** The TCRs from the most expanded clones in each organ at day 9 (a) or day 56 (c) was reconstructed and expressed in TCR-deficient murine T cell hybridomas and co-cultured with splenic DCs that had been pulsed with defined antigens for 24 hours. Left number plot represents clone distribution across organs for each clonotype/TCR, middle fraction plot represents cluster distriubtion for each clonotype/TCR and right plot represents antigen specificity for each clonotype/TCR. Data from 1 -2 independent experiments. TRACK mice were infected with influenza PR8 virus and CD4 T cells were enriched at day 9 (b) or day 56 (d) from each organ and stained with CFSE. The CD4 T cells were seperetaly co-cultured with splenic DCs that had been pulsed with indicated viral protein for 48 hours. CD4^+^Tomato^+^ T cells from each organ were analyzed for CD69 expression as a measurement of T cell activation. Representa­tive plots from 3 independent experiments, n = 3-4/group. Statistical significance were determined by one-way analysis of variance (ANOVA) with Tukey’s multiple-comparison test.

To confirm whether there is an organ-specific selection of TCR specificity across organs, we infected TRACK mice with influenza PR8, isolated CD4^+^ T cells from each organ, and co-cultured them separately with DCs that had been pulsed with influenza proteins. Consistent with our hybridoma data, influenza-specific CD4 T cells in the spleen had a significantly higher specificity for HA and neuraminidase (NA) than those found in the lungs (Fig. 6b). T cells residing in the medLN had intermediate specificity to HA and NA compared to those in the spleen and lungs. Similar dynamics were observed in terms of NP specificity across organs. When we tested the specificity to non-structural protein 1 (NS1), a protein highly expressed in infected cells, we found significantly larger specificity within the lungs than in T cells located in the medLN or spleen. We observed similar results when we pulsed our cocultures with viral particles (Extended Data Fig. 4f). Next, we explored the potential relationship between TCR usage and the emergence of specific T cell states during the effector stage of the immune response. To achieve this, we utilized the ‘clonotype bias’ metric to quantify whether individual clones are biased toward a specific subtype^37^. A clonotype bias of 1 signifies that all cells within a clonotype originate from the same subtype, whereas a clonotype bias of zero indicates that the clonotype precisely reflects the background subtype distribution of the entire sample. Our analysis demonstrates that most clonotypes were predominantly unbiased, resulting in the emergence of various CD4^+^ T cell states (Extended Data Fig. 4g).

At the memory stage, we observed increased clonal overlap across organs, in contrast to the compartmentalized repertoires observed during the effector phase (Fig. 6c). Of the 15 TCRs cloned at this stage, 14 showed productive expression (Extended Data Fig. 4d). Notably, although most memory-stage clones remained reactive to viral antigens (Fig. 6c and Extended Data Fig. 4e), their specificity was skewed toward viral proteins, such as NP and NA, suggesting a shift in antigenic dominance over time. Furthermore, the cluster distribution at this stage correlated with organ localization: Trm-like and cytotoxic clusters 7 and 8 were confined to the lungs, Tfh/Tsc-like cluster 2 was enriched in the medLN, and Tcm/Tsc-like cluster 5 was dominant in the spleen. To assess whether TCR specificity converges across organs, we infected TRACK mice with influenza PR8, isolated CD4⁺ T cells from each organ, and co-cultured them separately with DCs pulsed with influenza protein. Although influenza-specific CD4⁺ T cells in the lungs showed significantly greater reactivity to HA and NA than those in the spleen, the overall TCR specificity appeared to be normalized across organs (Fig. 6d). Consistent with our findings at the effector stage, we observed minimal clonotype bias during the memory phase (Extended Data Fig. 4h). Together, these data indicate that while organ-specific antigen presentation and recognition drive initial clonal expansion, long-term memory formation involves broader clonal dissemination and phenotypic adaptation to distinct tissue microenvironments. This convergence in antigen specificity may reflect ongoing antigen exposure across tissues or the preferential persistence of clones targeting specific viral epitopes.

## TCR functional avidity and organ antigenic landscape distinctly shape clonal dynamics

To assess the role of TCR affinity in clonal dynamics, including clonal expansion and distribution, we infected TRACK mice with influenza PR8 and isolated NP311–325-specific CD4⁺ T cells from the lungs, medLN, and spleen 9 days post-infection, followed by scTCR-seq analysis. TCRs from the most expanded clones across these organs were reconstructed and expressed in T cell hybridomas (Fig. 7a). The functional avidity of the selected TCRs was determined by stimulating the hybridomas with graded concentrations of their cognate NP311–325 peptide and calculating the EC50 for each TCR (Fig. 7b,c and Extended Data Fig. 4g). Because lower EC50 values indicate higher functional avidity, we found that higher-avidity TCRs were associated with larger clonal expansions. However, TCR avidity did not explain clonal distribution, as determined by the Gini index (Fig. 7d,e).

**Figure 7:**
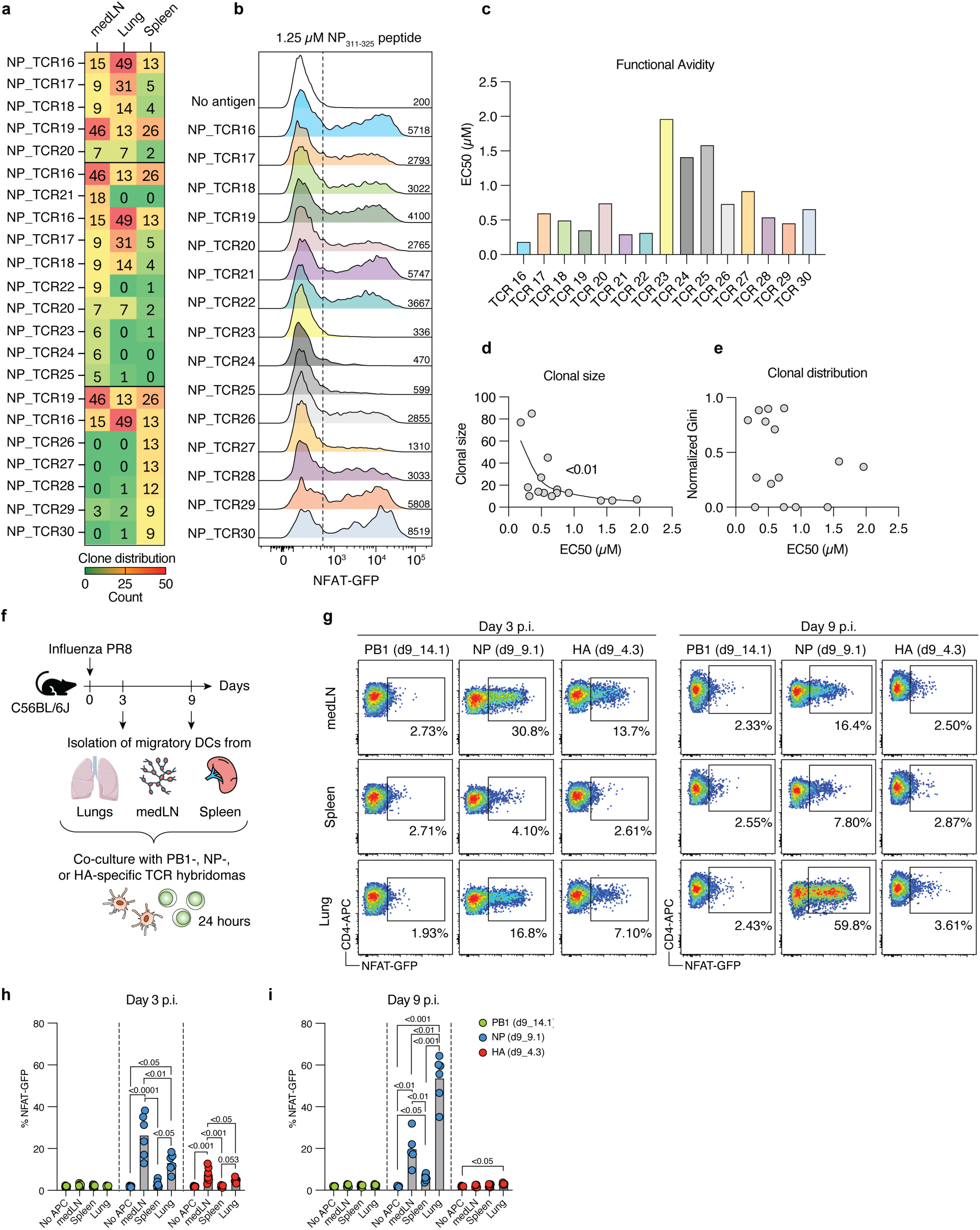
D**i**stinct contributions of TCR avidity and specificity to clonal dynamics a, NP_311 325_-specific T cells were isolated as single cells 9 days post influenza PR8 infection from the lungs, medLN ans spleen and their TCRs were seqenced. The TCRs from the top 5 expanded clones across each organ were expressed in NFAT-GFP T cell hybridomas and their functional avidity (EC50) were determined by NP311-325 peptide dilution experiment (b and c). d, Clonal size was correlated with functional avidity (EC50). e, Clonal distribution was correlated with functional avidity (EC50). Spearman’s rank correlation was used to assess the significance of the relationship between variables for d and e. f and g, C56BL6/J mice were infected with influenza PR8 and CD11c^int^MHCII^HI^ migratory DCs were isolated from the lungs, medLN and spleen at day 3 (h) or day 9 (i) post infection. The DCs were directly co-cultured with either PB1-, NP-, or HA-specific hybridomas for 24 hours. Pooled data from 2 independent experiments. Statistical significance were determined by one-way analysis of variance (ANOVA) with Tukey’s multiple-comparison test.

To investigate whether the antigen landscape varies across organs and over time, we infected C57BL/6J mice with influenza PR8 and isolated CD11c⁺MHCII^hi^ mDCs from the lungs, medLN, and spleen. These DCs were co-cultured with PB1-, NP-, or HA-specific T cell hybridomas without the addition of exogenous antigens to assess antigen presentation (Fig. 7f). At 3 days post-infection, DCs from the lungs and medLN presented NP antigens that induced comparable NFAT-GFP activation in NP-specific hybridomas, while only 2 of 6 mice had splenic DCs capable of inducing detectable NFAT expression (Fig. 7g,h). Additionally, DCs from the lungs and medLN presented HA antigens that elicited lower but significant NFAT responses in HA-specific hybridomas. Nine days post-infection, lung DCs presented NP antigens that triggered higher NFAT activation in NP-specific hybridomas, whereas medLN DCs maintained similar levels of NP presentation as on day 3 (Fig. 7i). HA presentation at day 9 was detected at low levels in the lung but remained significantly above the background level, whereas medLN and spleen DCs showed no detectable HA reactivity. PB1 presentation was not detected at any time point or in any of the organs. These data suggest that TCR functional avidity governs clonal expansion rather than distribution, and that dynamic, organ- and time-dependent antigen availability and presentation are major contributors to clonal selection during influenza infection.

## Discussion

Our findings provide critical insights into how tissue-specific conditions influence CD4⁺ T cell differentiation and clonal selection during infections. Previous studies have comprehensively defined the role of secondary lymphoid tissues in shaping T cell responses; our data extend this body of knowledge by revealing distinct organ-dependent transcriptional programs and clonal repertoire acquired by pathogen-specific T cells.

The observation that draining LNs preferentially support Tfh differentiation is consistent with their recognized role in facilitating B cell help in germinal center responses^38^. Additionally, our analysis implicating the spleen in the early dissemination of flu-specific CD4⁺ T cells to the lungs aligns with earlier findings suggesting a role for the spleen in generating systemically migratory and long-lived memory CD8^+^ T cells^27,29^. Mechanistically, it was suggested that migratory DCs from influenza-infected lungs migrate to the draining LNs first and continue their migration to the spleen, where they are found within 48 h of influenza infection. The high expression of the transcription factors Tcf7 and Klf2 by fate-mapped CD4^+^ T cells in the spleen early after infection not only marks their stem-like and migratory properties but may also explain their reduced Tfh differentiation^39–41^. This reinforces a model in which the spleen serves as an amplifier and distributor of systemic immune responses, complementing the local immune responses orchestrated by the draining lymph nodes. However, our data cannot directly distinguish whether the observed functional biases arise from activation and differentiation within the tissue microenvironment or from functional adaptation following migration across organs.

Our approach also enabled a broader assessment of the interplay between the T cell repertoire and pathogen-derived antigens during the immune response. Using TCR sequences as clonal barcodes, we observed that clonal sharing between the lungs and spleen was greater than that between the lungs and medLNs during the effector phase. One plausible explanation is that the induction of the migratory Tsc cell phenotype in the spleen directly leads to increased migration and distribution to the lungs during the effector phase. Mathematical modeling of influenza-specific CD8⁺ T cell responses also support this possibility by identifying the spleen as a key reservoir and source of effector T cells during infection^42^. This model suggests that the immune response is constrained more by the antigen presentation context than by the viral load in different tissues. These findings support a model in which the medLN is primarily responsible for initial priming, whereas the spleen functions as a central amplifier, expanding and systemically disseminating effector clones, including the lungs, as indicated by the splenectomy of TRACK mice during infection.

Furthermore, our findings on organ-specific T cell recognition patterns resonate with current concepts regarding antigen distribution dynamics dictated by pathogen and APC biology^43–46^. The influenza virus replicates intracellularly within lung epithelial cells, creating a distinct antigenic environment enriched for intracellular antigens involved in virus replication, such as polymerase PB1, which is a likely explanation for the T cell recognition biases towards this low-abundance protein in the lungs. In contrast, the early transport of antigens by migratory DCs to lymphoid organs and the spleen predominantly involves structural proteins such as HA and NP, reinforcing earlier predictions of how antigen transport may dictate the T cell repertoire^27,43^. Such patterns might differ significantly for extracellular pathogens or infections with distinct replication niches, affecting T cell priming and clonal selection in other contexts. Consistent with this, our ex vivo analysis of DC antigen presentation using hybridoma co-cultures revealed dynamic and organ-specific changes in antigen availability over time, further supporting the notion that tissue-resident APCs shape the evolving T cell landscape during infection.

The convergence of antigen specificity across tissues during memory, predominantly targeting viral proteins NA and NP aligns with theoretical models proposing affinity-based clonal competition and persistence driven by intermittent antigen exposure or homeostatic proliferation^47^. Our data also extend recent insights into retrograde migration, highlighting an increased clonal overlap between peripheral tissues and draining lymph nodes during the memory phase via the active migration of memory T cells^32,34^. Nevertheless, we cannot exclude the possibility that relative enrichment of specific clones within individual tissues contributes to this convergence independently of egress. In contrast, the clonal overlap between the lung and spleen remained static during the effector and memory phases. Although it is currently unclear whether T cells that exit the lung and migrate to draining LNs can continue to other sites, these observations point to a low-level continuous seeding of clones from the lung or medLN into the circulation and spleen. A similar phenomenon was recently reported for ex-Trm CD8⁺ cells during LCMV infection in a P14 TCR transgenic model^48^, further supporting the idea that memory T cell dynamics involve ongoing redistribution between tissues and lymphoid sites via circulation.

We did not observe clonal bias among the expanded clonotypes during either the effector or memory phases of influenza infection, supporting the “one clone, multiple fates” model of CD4⁺ T cell differentiation^37,49–51^. However, recent studies suggest that CD4⁺ T cell fate and memory potential can be influenced by TCR usage, peptide affinity, and the nature of the infection^49,52–54^. In particular, TCR affinity has been shown to not only affect memory formation but also determine the duration of T cell engagement with APCs within secondary lymphoid organs^55,56^. Our analysis of NP-specific TCRs suggests that whereas higher-avidity TCRs preferentially expand, antigen availability and local cues play a major role in the spatial segregation of T cells. Our results add a new layer to current models by showing that antigen distribution across tissues, in addition to local environmental cues and TCR affinity, likely plays a critical role in shaping T cell clonal selection, differentiation and memory potential.

Our study also establishes the TRACK system as a genetic tool for tracing recently activated T cells across organs during infection, capturing the earliest stages of de novo T cell activation. This system complements the recently described AgRSR model^57^, but with key distinctions: while AgRSR labels all recently TCR-activated T cells, TRACK selectively labels those that retain CD62L expression, thereby enriching for naïve-derived cells undergoing de novo activation.

Our findings highlight that tissue-specific priming environments and pathogen-dependent antigen distribution critically influence T cell differentiation, clonal selection, and memory formation. These insights could inform strategies to improve vaccination efficacy and manage infections by considering tissue-specific immune dynamics and pathogen biology.

## Limitations of the study

The TRACK model achieved approximately 35% labeling efficiency, leaving a substantial fraction of activated T cells untracked, which potentially influences the interpretation of clonal dynamics. Additionally, the *in vitro* assays used to confirm antigen specificity did not account for all labeled T cells. We also found that IFN-β can result in low levels of labeling in non-activated T cells, raising questions about possible bystander activation or alternative activation mechanisms not captured in these assays. Furthermore, our conclusions are currently limited to influenza virus infection; thus, the applicability of these conclusions to other pathogens with different replication niches or chronic infections and autoimmune conditions remains to be validated. Future studies leveraging the TRACK model across diverse infectious and inflammatory contexts will help address these limitations and refine our understanding of tissue-specific immune dynamics in humans.

## Acknowledgement

We are grateful to A. Rogoz and J. Bortolatto for embryo bioengineering, animal care, mouse colony management, and genotyping, RU Genomics core for assistance with sequencing, K. Gordon, and J.P. Truman for sorting, Yasmeen Khan and additional Rockefeller University employees for continuous assistance. We thank the NIH Tetramer Facility for providing the LLO and NP tetramers. We thank all the members of the Mucida, Victora (Rockefeller), and Lafaille (NYU) labs for fruitful discussions. This work was supported by The Swedish-American Foundation and The Swedish Research Council (R.P.), grant #2023/02507-7, São Paulo Research Foundation – FAPESP (H. A.), a Life Sciences Research Foundation postdoctoral fellowship (T.B.R.C), G.L.R. was supported by a scholarship from Fundação Maria Emília, FME (Brazil), and NIH grants R01DK093674 and R01DK113375, NIAID grant P01AI179273, the CZI Science, Food Allergy FARE/FASI Consortium, Stavros Niarchos Foundation Institute for Global Infectious Diseases, and The Howard Hughes Medical Institute (D.M.).

## Author contributions

Conceptualization: R.P. and D.M.; Investigation: R.P., H.A., T.C., G.L., A.S., and A.B.; Writing – original draft:

R.P. and D.M.; Editing: All authors; Funding acquisition: R.P. and D.M. Resources: T.C. and H.H.; Supervision: D.M.

## Methods Mice

Cd69-CreERT^2^ and Sell-DreERT^2^ mice were generated by CRISPR gene targeting in fertilized C57BL/6J (Jackson Laboratories) mouse embryos. Genetically engineered mice were backcrossed for 3-5 generations with C57BL/6J mice to remove off-target Cas9 genome modifications. TRACK mice were generated by intercrossing Cd69-CreER^T2^, Sell-DreER^T2,^ and Rosa26-CAG-RoxSTOPRox-LopxSTOPLoxp-tdTomato mice (Ai66). Ai66 mice were kindly provided by Hongkui Zeng (Allen Institute for Brain Science). B6.NPFlu (NP-specific TCR transgenic) mice were kindly provided by Susan Swain (U Mass Chan)^58^. The experimental mice were of the genotype Cd69^CreER/wt^ × Sell^DreER/wt^ × Rosa26^RCRL-tdT/RCRL-tdT^. Adult male and female mice (6–13 weeks old) were used in the experiments. Mice were housed at 22.2 °C and 30–70 % humidity in a 12 hours light/dark cycle with ad libitum access to food and water. All protocols were approved by the Institutional Animal Care and Use Committee of Rockefeller University (protocol number 22071-H).

## Plasmids

The Cd69-CreERT^2^ targeting vector was a modification of the pUC19 plasmid (Supplementary Fig. 1a). The stop codon was removed from the last exon (exon 5) of the Cd69 gene, and a P2A sequence was inserted immediately after the modified exon 5 sequence. The CreERT^2^ sequence was inserted directly after P2A. The targeting sequence was flanked by two homology arms (HArm) surrounding exon 5 at the *CdCS* gene locus: a long 2 kb arm at the 5’ end and a shorter 1 kb arm at the 3’ end. The homology arms were amplified by PCR from purified C57BL/6J hematopoietic stem cell DNA using primers that flanked the gene region. The 5’ HA end primer included the modified exon 5 without a stop codon. P2A and CreERT2 sequences were synthesized (IDT DNA). The pUC19 plasmid was linearized with XhoI (New England Biolabs), and the targeting vector was assembled with linear pUC19, P2A-CreERT^2^, 5’HArm, and 3’HArm using the NEB HiFi Assembly Kit (New England Biolabs). The Sell-CreERT^2^ targeting vector was a modified pUC19 plasmid (Supplementary Fig. 1b). The stop codon was removed from the last exon (exon 8) of the *Sell* gene, and a P2A sequence was inserted immediately after the modified exon 8 sequence. The DreERT^2^ sequence was inserted directly after P2A. The targeting sequence was flanked by two homology arms surrounding exon 8 at the *Sell* gene locus: a long 2.8 kb arm at the 5’ end and a shorter 1 kb arm at the 3’ end. Similarly, the targeting vector plasmid was assembled using the NEB HiFi Assembly Kit.

## Preparation of CRISPR injection mixes

Briefly, following the Easi-CRISPR protocol^59^, CRISPR guide RNAs were designed using CRISPR.mit.edu or CHOPCHOP. The protospacer sequence for the *CdCS* single guide (sg)RNA was AGCAAGCCCTCCAGATGACG (IDT DNA), and the protospacer sequence for *Sell* sgRNA was TCACAAAGGATGAATCAGTA (IDT DNA). The final concentrations of the components for the injection mixture were 50 ng/µl sgRNA, 50 ng/µl Cas9 protein (1081058, IDT DNA), and 15 ng/µl target vector plasmid in 80 µl ultrapure water (Gibco). The final injection mixes were passed through Millipore Centrifugal Filter units (UFC30VV25, EMD Millipore) and spun at 21,000 × g for 5 min, at room temperature.

## Microinjection of one-cell embryos

C57BL/6J mice aged 3–4 weeks were superovulated by intraperitoneal injection of 5 IU pregnant mare serum gonadotropin (hCG), followed by injection of 5 IU human chorionic gonadotropin 48 h later (Calbiochem, Millipore). Mouse zygotes were obtained by mating C57BL/6J stud males with superovulated C57BL/6 females. Fertilized mouse embryos were injected with the Cas9 protein, sgRNA, and target vector plasmid. Microinjection and mouse transgenesis experiments were performed as previously described^60^.

## Infections and treatments

Tamoxifen (Sigma) was dissolved in corn oil (Sigma) and 10 % ethanol, shaken at 45°C for 2-4 h, and stored at -20°C until use. TRACK mice were treated with four doses of Tamoxifen (5 mg/dose). Tamoxifen was administered via 100 µl oral gavage at 50 mg/mL on days -1, 1, 3, and 5 relative to the infection. To label circulating immune cells, mice were injected intravenously with 2 µg of anti-mouse CD45-FITC in 100 µl of phosphate-buffered saline (PBS), 3 min prior to euthanasia and lung tissue dissection. *L. monocytogenes* 10403S-inlA strain expressing full-length OVA (Lm-OVA) were grown overnight in brain heart infusion media. Mice were infected with 10^8^ colony-forming units (CFU) 24 h after oral treatment with 20 mg of streptomycin (Sigma-Aldrich) diluted in water. Influenza A/PR/8/34 virus (IAV PR8) stocks were provided by the Rice and Ravetch laboratories (Rockefeller University). For anesthesia, the mice were intraperitoneally injected with a mixture of ketamine (80 mg/kg; Zoetis, #54771-2013-1) and xylazine (8.8 mg/kg; Akorn, #07-808-1947). After mice were sufficiently anesthetized, 30μL of 25-30 pfu PR8 inoculum was applied to one nostril. The mice were monitored until they regained consciousness.

## Splenectomy

Mice were anesthetized with 2-5 % isoflurane and treated with Puralube eye ointment to prevent corneal drying. The abdominal flank fur was shaved, and the skin was sequentially disinfected with Betadine and 70 % ethanol. A small left subcostal incision was made to access the spleen. The spleen was gently exteriorized. The splenic vasculature and its connection to the pancreas were carefully cauterized using a heat pen, and the spleen was removed from the animal. The pancreas was repositioned within the abdominal cavity. The peritoneum was closed with absorbable Vicryl 4-0 sutures, and the skin was closed using clip sutures. Ketamine was administered for perioperative analgesia.

## Flow cytometry and staining strategy

Following antibodies was used: anti-mouse TCRβ-BUV395 and TCRβ-PE (clone: H57-597, eBioscience), anti-mouse CD4-BUV496 and CD4-APC (clone: GK1.5, BD), anti-mouse CD8β-BUV563 (clone: 53-6.7, BD), anti-mouse CD44-APC and CD44-A700 (clone: IM7, Biolegend), anti-mouse CD62L-BV605 (clone: MEL-14, Biolegend), anti-mouse CD45-FITC and CD45-PE/Cy7 (clone: 30-F11, Biolegend), anti-mouse PD1-APC/Cy7 (clone: J43, Biolegend), anti-mouse CD69-PerCP/Cy5.5 (clone: H1.2F3, eBioscience), anti-mouse CXCR5-BV650 (clone: L138D7, Biolegend). For live dead discrimination, we used Zombie Aqua or Zombie NIR (Biolegend). The following gating strategy was used to examine CD4^+^ T cells: single live lymphocytes (based on size and live/dead fixable dye Aqua or NIR stain), CD45iv^–^, CD45^+^, TCRβ^+^, CD8β^–^, and CD4^+^. For the analysis of fate-mapped T cells in TRACK mice, the following gating strategy was added: CD44^+^ and Tomato^+^. For sorting T cells subjected to DC co-cultures, we gated on single live lymphocytes CD45iv^–^, CD45^+^, CD8β^–^, CD44^+^, and Tomato^+^. Flow cytometry data were acquired on a Symphony A5 flow cytometer (Becton Dickinson, USA) and analyzed using FlowJo 10 software package (Becton Dickinson, USA). For tetramer staining, LLO₁₉₀₋₂₀₁ and NP₃₁₁₋₃₂₅ tetramers were obtained from the NIH Tetramer Core, and T cells were stained in cRPMI supplemented with 50 nM dasatinib for 1 h at 37°C and 5 % CO2.

## Cell culture

Fate-mapped CD44^hi^Tomato^+^ T cells were isolated using a FACS Symphony S6 (BD) cell sorter and stained with 5 µM CFSE (Sigma) following the manufacturer’s instructions. The CFSE-stained T cells were then co-cultured with antigen-pulsed DCs at a ratio of 1:5 (T cell:DC) for 48 h in 200 µL of RPMI 1640 (Gibco) supplemented with 10 % FBS (Sigma), 1 % Pen/Strep (Gibco), 1 % L-glutamine (Gibco), 1% Sodium Pyruvate (Gibco), 2 % Non-essential Amino Acids (Gibco), 2.5 % 1M HEPES (Gibco), and 50 μM 2-Mercaptoethanol (Sigma) in a 96-well round bottom plate (Corning) at 37°C and 5 % CO2. Splenic DCs were pulsed with either Influenza PR8 or heat-killed *L. monocytogenes* 10403S. The Proliferation Index was calculated using FlowJo as the total number of divisions divided by the number of cells that entered division. The percentage of the initial population was computed as the fraction of divided cells relative to the estimated initial cell count at the start of the culture. For some experiments, CD69 expression was used to determine the T cell activation. T cell and DC co-cultures were similar as above however, the DCs were stimulated with influenza PR8 or HA (provided by Krammer lab, Mount Sinai), NP, NS1 or M1 (SinoBiological) at 1 µg/well. For the isolation of dendritic cells, C57BL/6J spleens were dissected into cold RPMI 1640 supplemented with 10 % heat-inactivated fetal bovine serum (Hyclone), 2 mM L-glutamine, 100 U/ml of penicillin, 100 μg/ml of streptomycin sulfate, 1 mM sodium pyruvate, 0.1 mM non-essential amino acids, 10 mM HEPES (all from Gibco), and 50 μM β mercaptoethanol (Sigma). Spleens were finely chopped and incubated in 400 U/ml Collagenase D (Roche) in supplemented RPMI for 25 min at 37°C and 5 % CO2. Single-cell suspensions were extracted from the connective tissue by taking up and resuspending the digests five times. Dendritic cells were enriched using anti-CD11c magnetic beads (Miltenyi Biotec).

## Lymphocyte isolation

Lymphocytes from the lungs were isolated by digestion the tissue in 3 ml RPMI 1640 supplemeted with 2% FBS (Hyclone), 10 mM HEPES (Gibco), 100 μg/ml DNaseI (Roche) and 1.5 mg/ml Collagense A (Roche) utilizing C Tubes (Miltenyi Biotec) with a gentleMACS Tissue Dissociator (Miltenyi Biotec) using the 37C_m_LDK_1 program. Mononuclear cells were isolated by gradient centrifugation using Percoll. Small intestine intraepithelial and lamina propria lymphocytes were isolated as previously described^15^. Briefly, the small intestines were harvested and washed in PBS and 1 mM dithiothreitol (DTT), followed by 30 mM EDTA. Intraepithelial cells were recovered from the supernatant of the DTT and EDTA washes. Lymphocytes from the lamina propria were obtained by digesting the tissue in 5 ml of RPMI, 2% FBS, 100 μg/ml DNaseI (Roche), and 2 mg/ml Collagenase 8 (Gibco) for 45 min at 37°C. Mononuclear cells were isolated by gradient centrifugation using Percoll. Lymphocytes from the spleen were isolated by mechanical dissociation and filtered through a 40 µm nylon filter. Erythrocytes were removed using RBC Lysis Buffer (Gibco) and washed with PBS (Gibco). Lymphocytes from the lymph nodes were isolated by mechanical dissociation and filtered through a 40 µm nylon filter. For CD4 T cell co-cultures, we used negative selection (biotinylated anti-CD8β, anti-CD11b, anti-CD11c, anti-Ter119, anti-NK1.1, anti-CD45R) with anti-biotin magnetic beads (Miltenyi Biotec) to enrich for CD4 T cells. TotalSeq-C antibodies (Biolegend) were used to hashtag CD4^+^CD44^hi^Tomato^+^ T cells isolated on BD Symphony S6 (BD) prior to droplet based scRNAseq. Following antibodies was used: Hashtag 1 C0301 (clone: M1/42, Biolegend, B381758, 1:100), Hashtag 2 C0302 (clone: M1/42, Biolegend, B364444, 1:100), Hashtag 3 C0303 (clone: M1/42, Biolegend, B344687, 1:100), Hashtag 4 C0304 (clone: M1/42, Biolegend, B383254, 1:100), Hashtag 5 C0305 (clone: M1/42, Biolegend, B348910, 1:100), Hashtag 6 C0306 (clone: M1/42, Biolegend, B357045, 1:100), Hashtag 7 C0307 (clone: M1/42, Biolegend, B364170, 1:100), Hashtag 8 C0308 (clone: M1/42, Biolegend, B341501, 1:100), Hashtag 9 C0309 (clone: M1/42, Biolegend, B344528, 1:100), Hashtag 10 C0310 (clone: M1/42, Biolegend, B352462, 1:100), Hashtag 11 C0311 (clone: M1/42, Biolegend, B389893, 1:100), Hashtag 12 C0312 (clone: M1/42, Biolegend, B384872, 1:100)

## Single cell RNA sequencing

Raw FASTQ sequencing files were processed using Cell Ranger v7.0.1 for both gene expression and feature barcode quantification. The resulting gene expression and antibody-derived tag (ADT) matrices were loaded into R and analyzed using the Seurat package (v5.3). ADT matrices were normalized using the centered log-ratio (CLR) method, and cell classification was performed using the HTODemux function. Cells exhibiting more than one ADT peak were identified as doublets and excluded from downstream analyses. Next, we removed cells showing more than 5% of mitochondrial genes from the analysis. The expression data were normalized using the SCTransform function. Dimensionality reduction was performed via principal component analysis (PCA), and UMAP was applied to visualize the clusters. Differentially expressed genes were determined using the Wilcoxon rank-sum test available with the FindMarkers function from the Seurat package. Multiple test correction was performed using the Bonferroni method, and genes containing less than an adjusted p-value of 0.05 were considered for downstream analysis. R-generated plots were produced using ggplot2 v3.5.2.

## Plate-based single cell TCR sequencing

Single cells were index-sorted using a FACS Symphony S6 into 96-well plates containing 5μL of lysis buffer (TCL buffer, QIAGEN 1031576) supplemented with 1 % β-mercaptoethanol and frozen at −80 °C prior to RT-PCR. RNA and RT-PCRs for TCRβ were prepared as previously described^61^. PCR products for TCRβ were multiplexed with barcodes and submitted for MiSeq sequencing using True Seq Nano kit (Illumina)^62^. Fastq files were de-multiplexed and paired-end reads were assembled at their overlapping region using the PANDASEQ^63^ and FASTAX toolkit. Demultiplexed and collapsed reads were assigned to the wells according to the barcodes. Fasta files from MiSeq sequences were then aligned and analyzed on IMGT (http://imgt.org/HighV-ǪUEST)^64^. Cells with identical TCRβ CDR3 nucleotide sequences were considered to be the same clones.

## T cell receptor and clonal repertoire analysis

Clonal TCR analysis was performed by processing the filtered_contig_annotations.csv output file from Cell Ranger. Briefly, only cells containing TRA, TRB, and double TRA_TRB contigs were used for statistical analysis. Cells harboring the same V and J genes and CDR3 length, as well as TRA and TRB pairing, were annotated as clonally related. Cells containing only TRA or TRB were also assigned as being part of the same TRA_TRB core if they had an exact CDR3 sequence. Circos plots were generated using Circos v0.69. Morisita horn overlap index was calculated with mh(data, CI = 0.95, resample = 100) function in the divo v1.0.2 R package. To measure cell state-TCR relationship, we used the ‘clonotype bias’ metric^37^: clonotype bias (cb) = maxi [(fi - qi)/ (1-qi)] where fi is the observed frequency for subtype i in the clonotype, and qi is the background frequency of subtype i in the whole sample. To assess the statistical significance of the measured cb scores, we generated N random binned permutations of the observed clonotype data, preserving clonal size (clone size > 5) and global subtype background frequencies. Z-scores for the observed clonotype bias scores were then calculated as the number of standard deviations from the background mean (Z-score > 1.96 corresponds to a p-value < 0.05).

## Generating TCR hybridomas

We used the retroviral vector pMSCV-mCD4-PIG TCR-OTII as previously described to transduce the NFAT-GFP 58α^−^β^−^ hybridoma cells with synthetic TCR constructs^65^. TCR sequences are listed in Supplementary Table 1. pMSCV-mCD4-PIG TCR-OTII, our backbone vector for TCR expression, was provided to Twist Bioscience. Sequences of the TCR α and β genes selected (30 TCRs in total) were obtained from the filtered_contig_annotations.csv output file from scRNA-seq. A target sequence consisting of the TCR α and β genes separated by a self-cleaving P2A peptide was synthesized (Twist Bioscience) and used to replace the OTII TCR gene in the pMSCV-mCD4-PIG plasmid. The Platinum-E retroviral packaging cell line (RV-101, Cell Biolabs) was used for the viral packaging of pMSCV-mCD4-PIG TCR vectors generated by Twist. Platinum-E cells were cultured in a 96-well plate to 60–80% confluence and transfected with pMSCV-mCD4-PIG TCR vectors as a DNA–lipid complex using Lipofectamine 3000 (L3000001, ThermoFisher Scientific) following the manufacturer’s instructions. The culture supernatant containing viral particles was harvested 24 h after transfection. Next, 2.5 x 10^4^ NFAT-GFP cells were centrifuged, resuspended in 200 μl of the virus-containing supernatant supplemented with 10 μg/ml protamine sulfate (MP Biomedicals), and plated in U-bottom 96-well plates. NFAT-GFP cells were spin-transduced by centrifugation at 1,000 × g at 32 °C for 120 min, pipetted up and down, and cultured overnight at 37 °C in a 5% CO 2 incubator. Cells were then transferred into a flask and incubated for 1–2 weeks with 10 ml of culture medium (DMEM with 10% FCS, penicillin-streptomycin, and 2 mM L-glutamine) under puromycin selection (1 μg/ml). After selection and propagation, TCR expression in NFAT-GFP cells was confirmed using flow cytometry.

## TCR hybridoma and DC co-cultures

First, 10^5^ splenic DCs were cultured in 100 µL in a 96-well U-bottom plate and pulsed with particles, molecules, or proteins of interest for 1-2 h. For PR8 virus particles, 10^5^ pfu/well was used. Proteins M1, NS1, NS2, HA, NA and NP (Sino Biological) was used at a concentration of 1 µg/well. For some experiments, NA and HA were purified and provided by the Krammer Laboratory at the Icahn School of Medicine at Mount Sinai. PB1 was expressed in the pET-29b(+) plasmid transduced into E. Coli. PB1 protein was produced and PB1 containing bacterial lysate were generated by 10 min heat shock in 0.5 ml deionized water, 0.5 µl lysate was used for DC pulsing. For the positive control, 0.5 µg of anti-mouse CD3ε (clone: 145-2C11, Biolegend) was used. Next, 2 x 10^4^ TCR-transduced NFAT-GFP cells were cocultured in a 96-well plate with the pre-pulsed DCs following co-culture at 37 °C in a 5 % CO 2 incubator. After 24 h of co-culture, cells were stained for Live/Dead Zombie NIR, TCRβ-PE, and CD4-APC. TCR stimulation was evaluated by measuring GFP expression by viable TCRβ^+^CD4^+^ hybridomas. Tissue derived TCRβ^−^, CD45R^−^, CD64^−^, NK1.1^−^, TCRγδ^−^, Ly6G^−^, CD11c^int^ and MHCII^hi^ mDCs were isolated from indicated tissues after digestion of with Collagenase (Collagenase A for lungs or Collagenase D for medLN or Spleen) by cell sorting on a FACS Symphony S6 (BD) and co-cultured with indicated TCR hybridomas in TCM media at a ratio of 1:1. Antibodies used for cell sorting of mDCs: TCRb-APC/Cy7 (clone: H57-597, Biolegend), CD45R-A700 (clone: RA3-6B2, Biolegend), NK1.1-PCP5.5 (clone: PK136, eBioscience), TCRgd-PCPeFlour710 (clone: eBioGL3, eBioscience), CD64-PCPeFlour710 (clone: X54-5/7.1, eBioscience), Ly6G-PCP (clone: 1A8, Biolegend), MHCII-BUV395 (clone: 2G9, BD), CD11c-BUV496 (clone: N418, BD), CD103-BV785 (clone: 2E7, Biolegend), CD8a-BUV805 (clone: 53-6.7, BD).

## Statistical analyses

Statistical analyses were performed using GraphPad Prism v.10. Flow cytometry analysis was performed using FlowJo software. Data in graphs show mean and p values <0.05 were considered to be significant. Adobe Illustrator was used to assemble and edit the figures. R Studio Version 2023.06.0+421 and R version 4.3.2 was used to analyze scRNAseq data.

---

**Extended Data Fig. 1:**
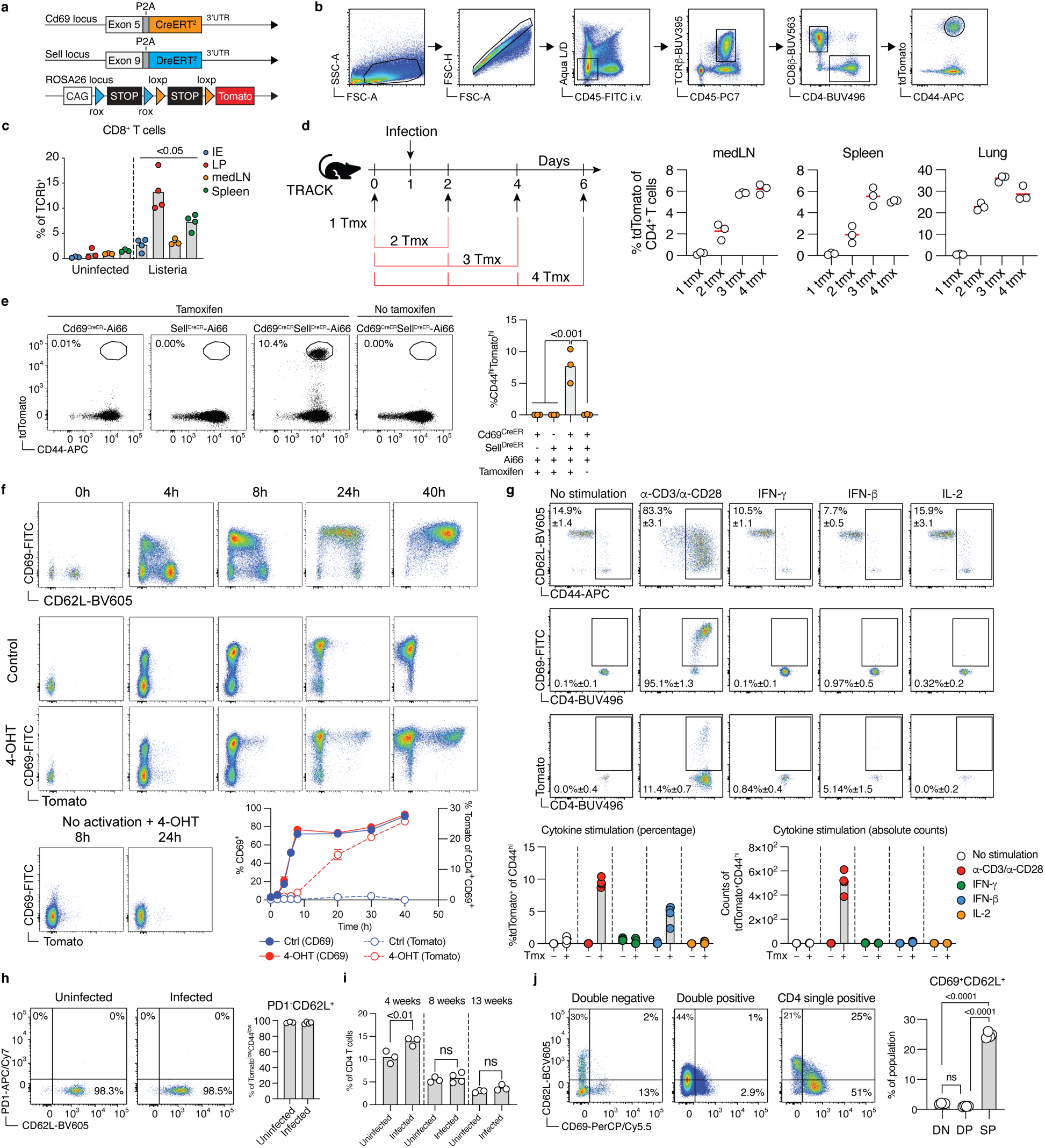
Tracking of pathogen-specific CD4^+^ T cells during intestinal and respiratory infection. a, Overview of genetic strategy regarding Cd69-CreER and Sell-DreER mice, b, Gating strategy for CD4^+^CD44^hi^Tomato^+^ T cells, c, TRACK mice were infected with L. monocytogenes and CD8^+^CD44^hi^Tomato^+^ T cells were analyzed at day 9 post infection. Representative plot. Minimum 3 independent experiments, n=3-5. Statistical signifi­cance was tested between infected organ to uninfected organ by Student’s T-test. d, TRACK mice were treated with tamoxifen as described and medLN, spleen and lung were analyzed for tdTomato labeling of CD4^+^CD44^hi^ T cells 9 days post influenza PR8 infection. n=12, data from one experiment, e, Stated mice were infected with L. monocytogenes and CD4^+^CD44^hi^Tomato^+^ were analyzed in the LP 9 day post infection. Representative plot from 2 independent experiments, n=1-2 per experiments. One-way analysis of variance (ANOVA) with Tukey’s multiple-comparison test, f, T cells from the spleen of TRACK mice were isolated and stimulated with anti-CD3/an-ti-CD28 with or without 4-Hydroxytamoxifen (4-OHT). CD69, CD62L and Tomato expression on CD4^+^ T cells were measured over time. Pooled data from two experiments, n = 6. g, Similar to e, but T cells were stimulated with IFN-y, IFN-p or IL-2 without CD3/CD28 stimulation. CD69, CD62L, CD44 and Tomato expression were measured on CD4^+^ T cells 24 hours post stimuilation. Data from 4 mice, h, i, Analysis of CD4^+^CD44^low^Tomato^+^ within the spleen at 4, 8 or 13 weeks of age. Representative plot of minimum 3 independent experiments. n=3-5, Student’s T-test. j, Analysis of CD69 and CD62L expression within the thymus in TRACK mice at 8 weeks of age. Pooled data from 2 independent experiments, n=6. One-way analysis of variance (ANOVA) with Tukey’s multiple-comparison test.

**Extended Data Fig. 2.**
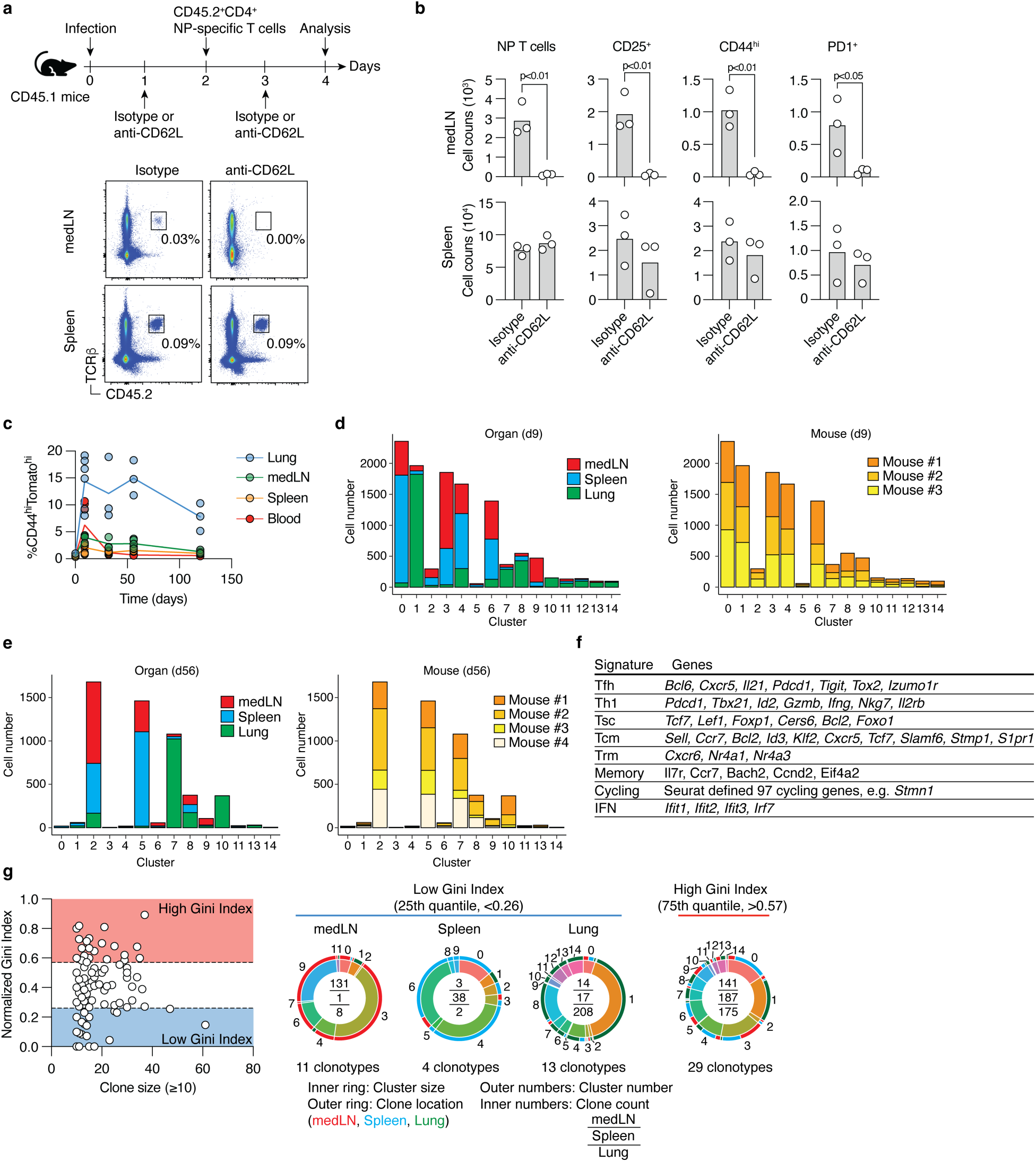
: Analysis of RNAseq data from influenza infected TRACK mice at day 9 and day 56 post infection. a and b, CD45.1 isogenic B6 mice were infected with influenza PR8 and treated with anti-CD62L at day 1 and 3. NP-specific naive 500.000 CD4 T cells were transferred to the reciepeints at day 2 post infection and the transferred T cells in the medLN and spleen were analyzed for activation markers at day 4 post inefection. c, Analysis of CD4^+^CD44^hi^Tomato^+^ T cells across timepoints and organs in TRACK mice post influenza PR8 virus infection. Pooled data from 5 independent experiments n=17. d, Cell counts across clusters divided by either organs (left) or mice (right) at day 9 post infection, n=3. e, Cell counts across clusters divided by either organs (left) or mice (right) at day 56 post infection, n=4. f, Gene signatures used for classifications in UMAP plots, g, All expanded clonotypes (clonal size >10) at day 9 post infection were analyzed to determine the Gini index per clonotype across medLN, spleen and lung. The top 25th and 75th quantile of the Gini index where determined and those selected clonotypes, i.e. low Gini index clonotypes (<0.26) or high Gini index clonotypes (>0.57) where pooled and analyzed for cluster distribution. Outer numbers: Cluster, Outer ring: organ origin, Inner ring: Cluster size, Inner number: Clone counts across organs. Pooled data from 3 mice.

**Extended Data Fig. 3.**
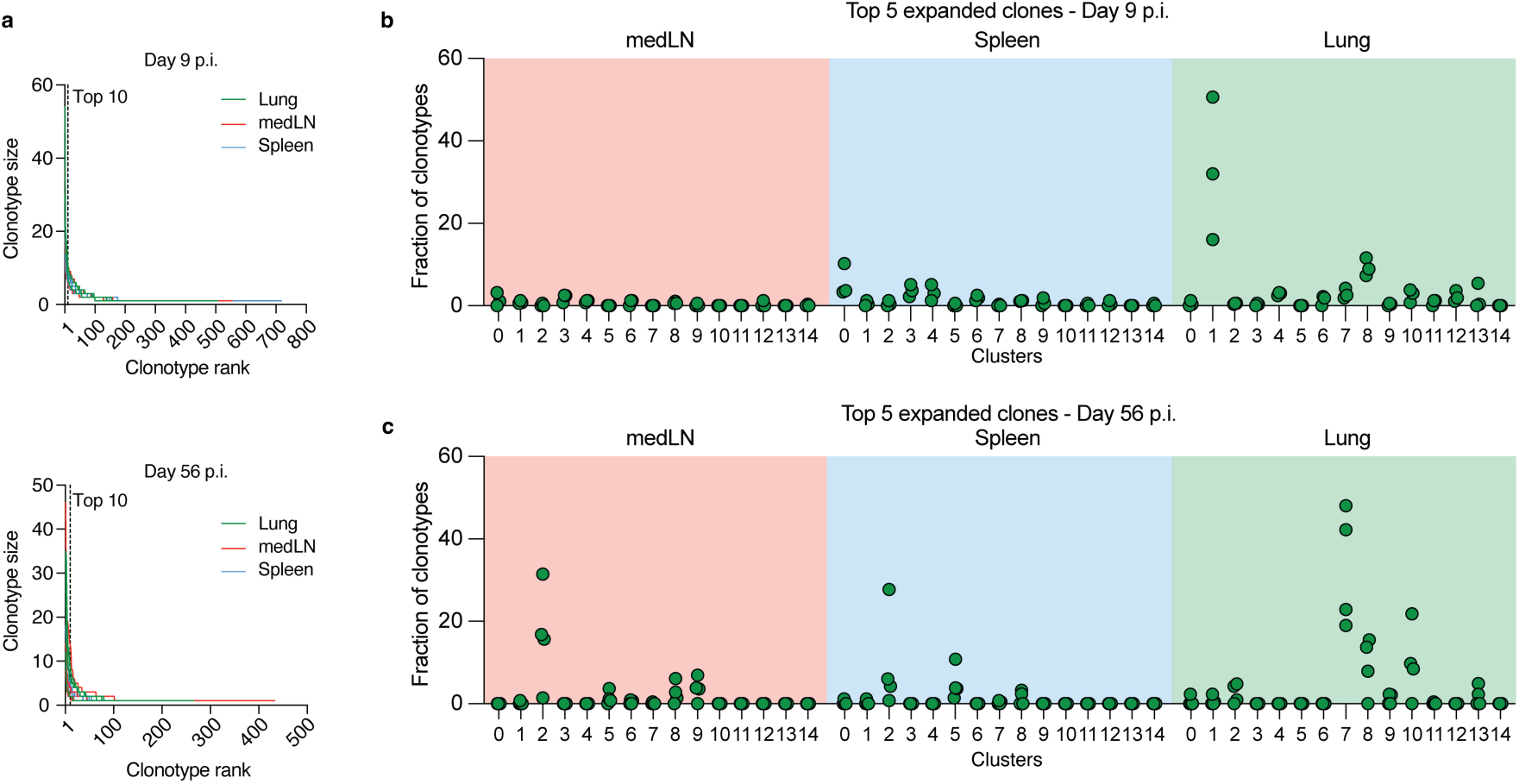
**: Distinct clonal expansion and distribution across organs.** a, Clonotype size ranking from scRNAseq data across organs and timepoints. Dashed x-axis line indicates top 10 expanded clones. Pool ranked clonotypes from 3 mice for d9 p.i. and from 4 mice for d56 p.i. b, Cluster and organ distribution of top 5 expanded clones located in the lungs at day 9 post infection. Each dot represents one unique mouse. n=3. c, Cluster and organ distribution of top 5 expanded clones located in the lungs at day 56 post infection. Each dot represents one unique mouse. n=4.

**Extended Data Fig. 4.**
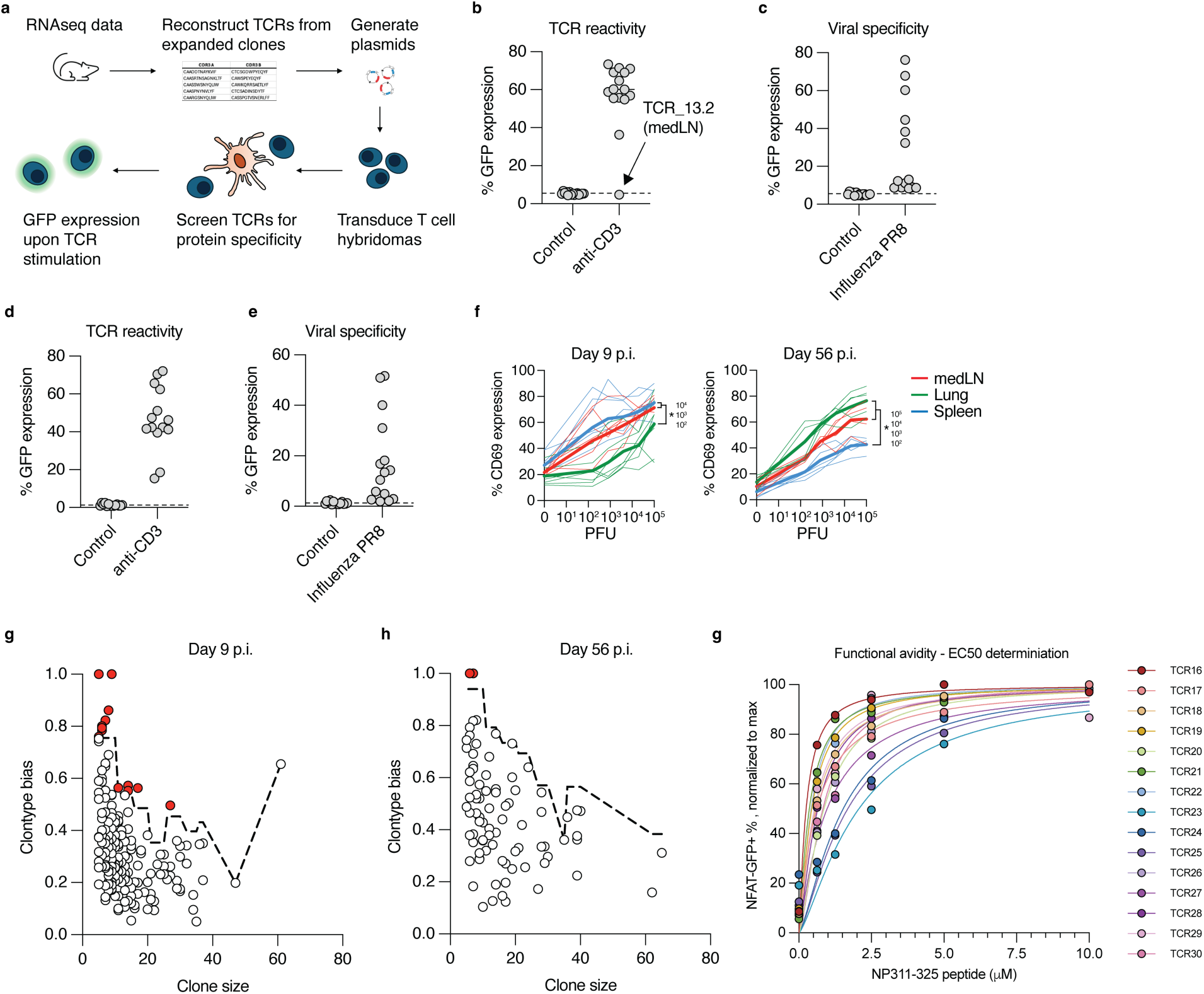
: Distinct clonal expansion and distribution across organs. a, Experimental overview of testing TCR specificty with murine T cell hybridomas. b, The 15 most expanded TCRs across lungs, medLN and spleen from day 9 post infection stimulated with anti-CD3 to determine functinality. TCRs are isloated from the scRNAseq data in figure 3. c, TCR hybridomas from day 9 were tested for viral specificty by co-culturing them with splenic DCs that had been pulsed with 10^5^ plaque forming units (pfu) of PR8 particles for 24 hours, d, The 15 most expanded TCRs across lungs, medLN and spleen from day 56 post infection, stimulated with anti-CD3 to determine functinality. TCRs are isloated from the scRNAseq data in figure 3. e, TCR hybridomas from day 56 were tested for viral specificty by co-culturing them with splenic DCs that had been pulsed with 10^5^ pfu of PR8 particles for 24 hours, f, TRACK mice were infected with influenza PR8 and CD4^+^ T cells were enriched from the lungs, medLN and spleen, 9 or 56 days post infection. The isolated CD4 T cells were stained with CFSE and co-cultured with splenic DCs that were pulsed with various amounts of PR8 viral particles for 48 hours. Pooled data from 2 independent experi­ments, n = 5-7/group. One-way analysis of variance (ANOVA) with Tukey’s multiple-comparison test, per dilution point. * = p < 0.05. g, Clonotype bias at day 9 post infection (p.i.), analysis only considered expanded clonotypes with 5 or more cells, and corrected for clonotype size by generating expected background distributions by random permutation (dotted line). Red circles indicate Z score > 1.97 and p < 0.05. Pooled RNAseq data from 3 individual mice, h, Clonotype bias at day 56 p.i., analysis only considered expanded clonotypes with 5 or more cells, and corrected for clonotype size by generating expected background distributions by random permutation (dotted line). Red circles indicate Z score > 1.97 and p < 0.05. Pooled RNAseq data from 4 individual mice, g, NP_311 325_-specific TCR hybridomas was co-cultured with splenic CD11c^+^ DCs and stimulated with various concentrations of NP311-325 peptide (10 pM, 5 ^M, 2.5 pM, 1.25 pM, 0.63 /jM and 0 pM) for 24 hours and NFAT-GFP expression was measured.

**Supplementary Fig. 1.**
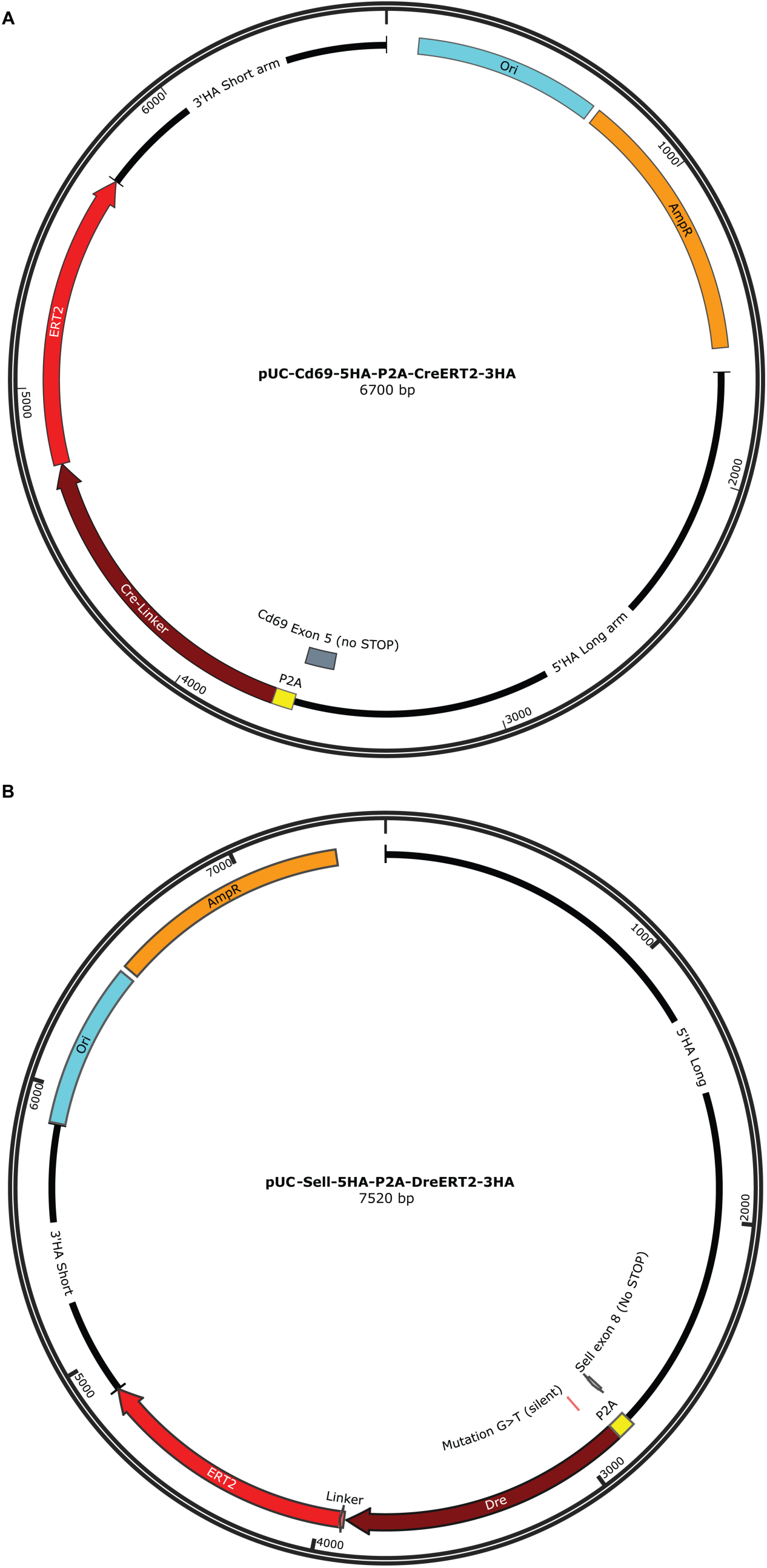
: Target vector plasmids for Cd69-CreERT^2^ and Sell-DreERT^2^ mice. (A) Cd69-CreERT2 plasmid (B) Sell-DreERT2 plasmid

**Supplementary table 1.**
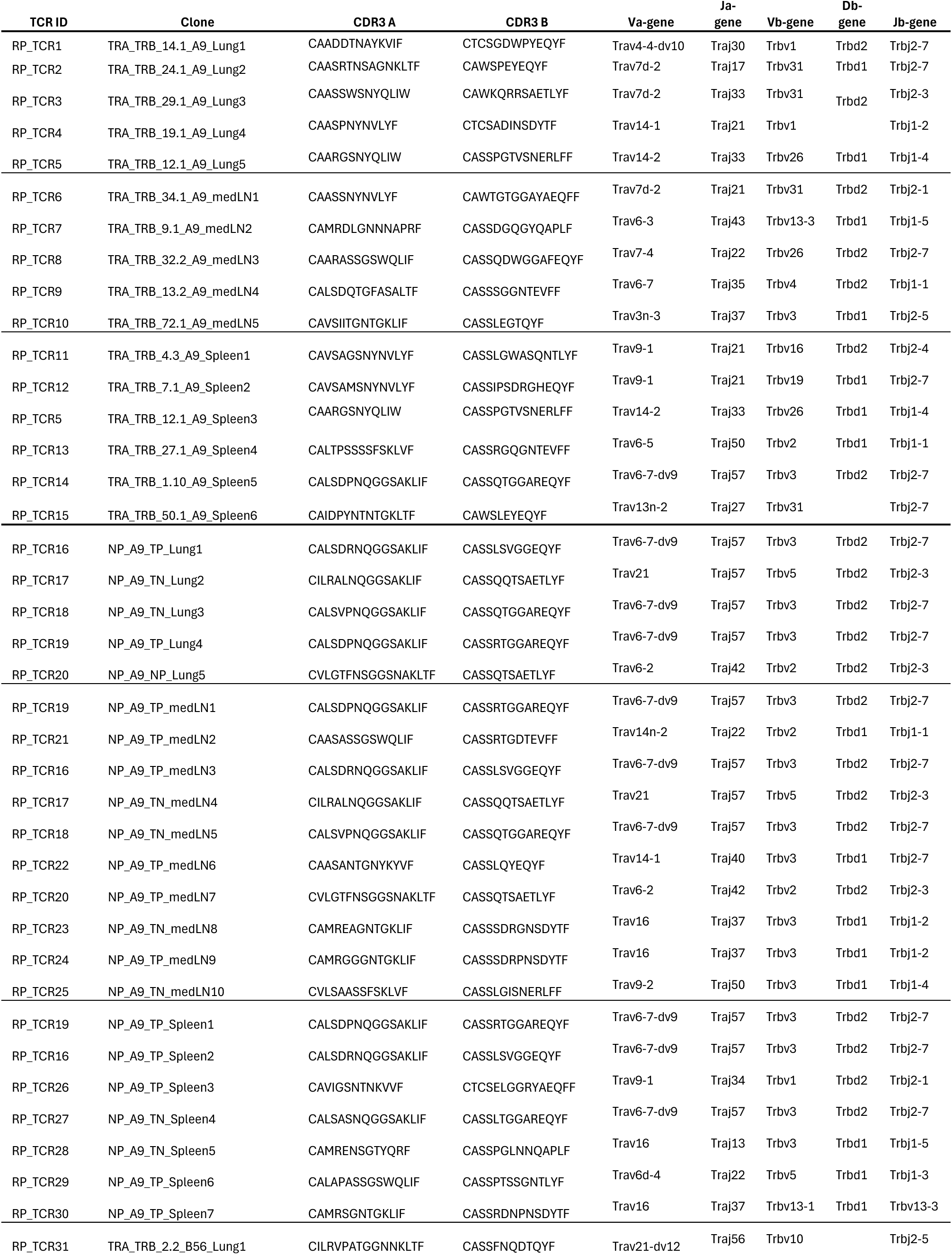
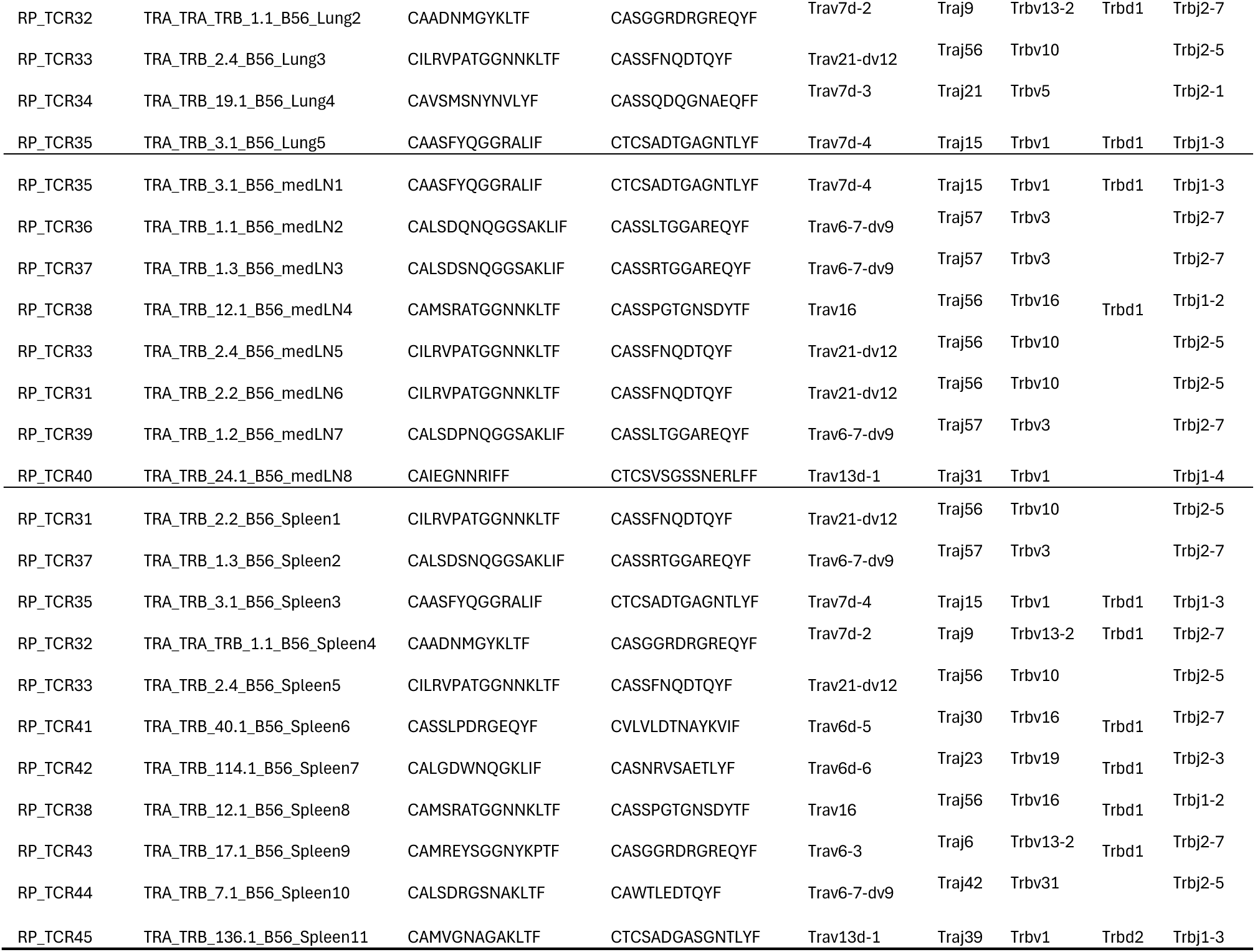
TCRs used for hybridomas, isolated from most expanded clones across organs

